# Context-dependent effects of IL-2 rewire immunity into distinct cellular circuits

**DOI:** 10.1101/2020.12.18.423431

**Authors:** Carly E. Whyte, Kailash Singh, Oliver T. Burton, Meryem Aloulou, Alena Moudra, Carlos P. Roca, Francisco J. Naranjo, Félix Lombard-Vadnais, Lubna Kouser, Tino Hochepied, Timotheus Y. F. Halim, Susan Schlenner, Sylvie Lesage, James Dooley, Adrian Liston

## Abstract

Interleukin 2 (IL-2) is a key homeostatic cytokine, with potential therapeutic applications in both immunogenic and tolerogenic immune modulation. Clinical application has been hampered by pleiotropic functionality and wide-spread receptor expression, with unexpected adverse events during trials. To characterize the IL-2 homeostatic network, we developed a novel mouse strain allowing IL-2 production to be diverted. Rewiring of IL-2 production to diverse leukocyte sources allowed the identification of contextual influences over IL-2 impact. Network analysis identified a priority access for Tregs, and a competitive fitness cost induced among both Tregs and conventional CD4 T cells for IL-2 production. CD8 T cells and NK cells, by contrast, exhibited a preference for autocrine IL-2 production. IL-2 sourced from dendritic cells amplified the Treg circuit, while IL-2 produced by B cells induced two context-dependent circuits: dramatic expansion of CD8^+^ Tregs and ILC2 cells. The former was associated with an unexpected concentration of rare CD8^+^ Tregs in B cell zones, while the latter drove a downstream, IL-5-mediated, eosinophilic circuit. The source-specific effects demonstrate the contextual influence of IL-2 function and potentially explain unexpected adverse effects observed during clinical trials of exogenous IL-2. Targeted IL-2 production therefore has the potential to amplify or quench particular circuits in the IL-2 network, based on clinical desirability.

**Graphical abstract:** 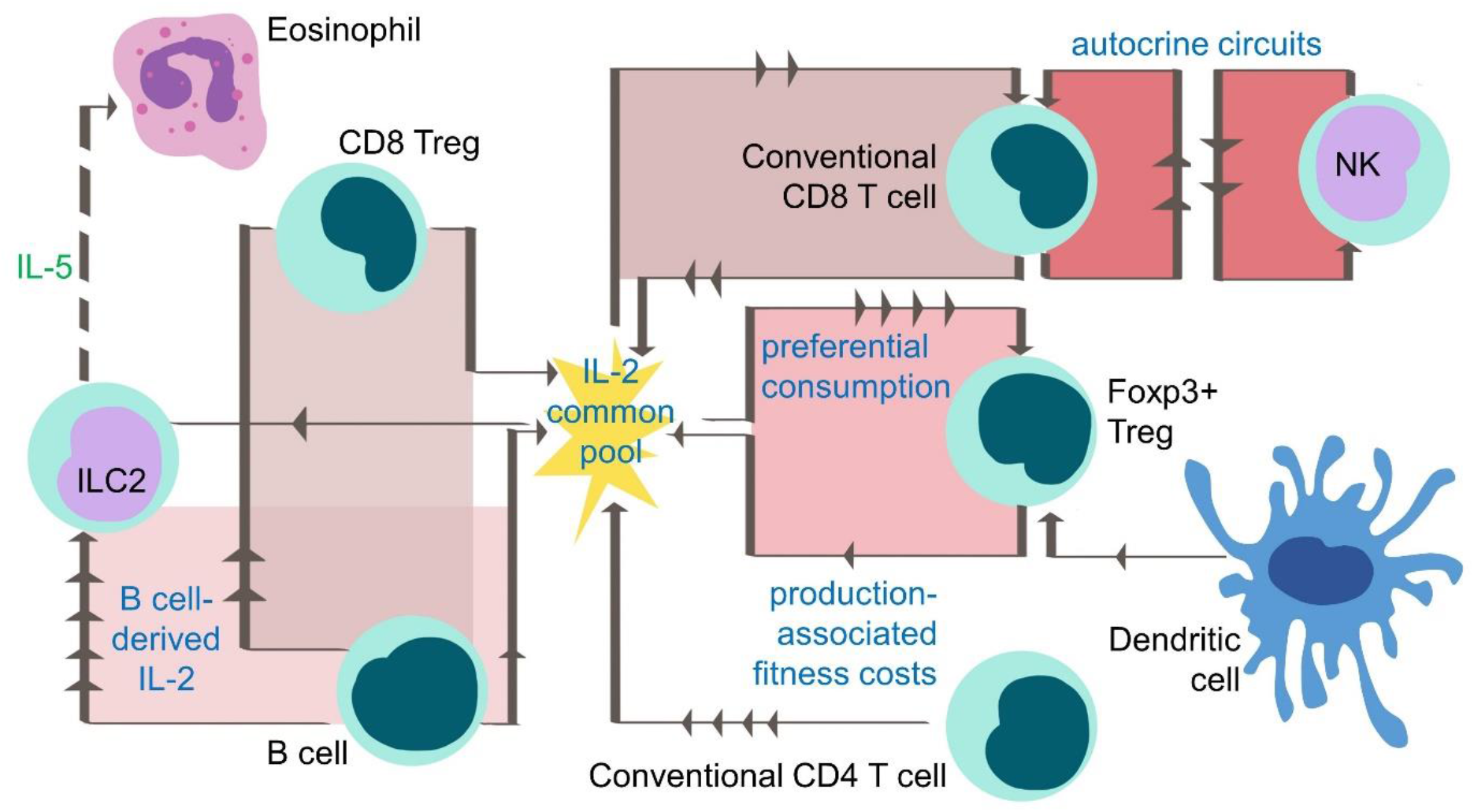

## Introduction

Interleukin 2 (IL-2) is one of the key homeostatic cytokines controlling the immune system (Malek, 2008). Among the first to be discovered, our understanding of IL-2 has shifted from a broad-utility T cell growth factor into the central maintenance factor for regulatory T cell (Treg) “fitness” (Fontenot et al., 2005). Individual cellular circuits of IL-2 signaling have been identified and characterized in depth. Arguably, the most important of these involves the early production of IL-2 by activated CD4 T cells (Amado et al., 2013), which in turn drives the expansion of Treg numbers via altering the proliferation and apoptotic kinetics (Obata et al., 2014; Pierson et al., 2013; Shi et al., 2018). While this creates a closed circuit, with negative feedback properties (Liston and Gray, 2014; Smith and Popmihajlov, 2008), this network can be rewired upon a change of context, such as driving a positive feedback loop of inflammatory CD8 T cell activation (Humblet-Baron et al., 2019; Humblet-Baron et al., 2016). Here the rewiring is based in part on the expression of the high affinity (K_d_ ≈ 10^−11^ M) trimeric receptor of CD25, CD122 and CD132, expressed constitutively by Tregs and upon activation in CD8 T cells. The preferential capture of IL-2 by CD25 provides a competitive advantage over cells (conventional CD4 T cells, naïve CD8 T cells and NK cells) that largely express the intermediate affinity (K_d_ ≈ 10^−9^ M) dimer of CD122 and CD132 (Spangler et al., 2015). The existence of alternative affinity receptors, dynamically regulated in quantity and expressed on multiple cell types, demonstrates the potential plethora of cellular circuits that could be controlled by IL-2.

A complete understanding of the network effects of IL-2 on immune homeostasis will need to go beyond the identification of pairwise circuits. Homeostatic networks can be defined based on controlled variables and regulated variables, with controllers acting on plants to stabilize the system (Kotas and Medzhitov, 2015). Using these definitions, activated CD4 T cells can be considered plants, producing the controlled variable IL-2 to regulate Treg numbers. However, in a biological system as complex as the immune system, the distinction between these homeostatic components is obscured by multiple layers of regulation, interconnection between variables, conditional dependence of signals and cell types simultaneously acting in multiple roles (Kotas and Medzhitov, 2015). Tregs, for example, can be considered both a regulated variable, stabilized by IL-2, and a controller which acts on CD4 T cells to tune IL-2 expression. The broader the set of cells and potential interactions considered, the more complex potential effects of each network component can be. For signaling components with high biological potency, such as IL-2, a more complete understanding of the network may depend upon top-down measurements of network-level perturbations, rather than bottom-up construction of the network from the building blocks of defined circuits.

Moving towards a more complete understanding of the IL-2 network does not solely serve as a proof-of-principle for dissecting the complex biology of pleiotropic cytokines. IL-2 is also an actively investigated therapeutic drug, the subject of hundreds of ongoing clinical trials. The importance of the contextual aspects of the IL-2 network lies in the diametrically opposed clinical uses. IL-2 has been adopted for its ability to distort immunity towards either an immunostimulatory or immunosuppressive state, based on the opposing targets of conventional T cells and Treg, respectively. Initially FDA-approved for treatment of metastatic renal cell carcinoma and melanoma in the 1990’s (Rosenberg, 2014), prior to the resurgence of Treg biology, high doses of IL-2 are used as stimulatory immunotherapy. The key target of this approach is generally anti-tumor CD8^+^ T cell responses, although trials are also underway to enhance infectious immunity (Pol et al., 2020). Conversely, the identification of CD25^+^ Tregs led to the design of low doses IL-2 trials, aimed at enhancing Treg numbers and function. These trials aim to suppress pathological and autoimmune responses, in diseases ranging from type I diabetes to graft-versus-host disease (Hartemann et al., 2013; Koreth et al., 2011; Matsuoka et al., 2013). Despite promising results in a proportion of patients, the therapeutic efficacy of IL-2 has been hampered by the pleiotropic effects on diverse cell types. Toxicity is often observed, with a vast array of side effects reported in patients, including vascular leak syndrome, hypotension and end-organ dysfunction, often leading to discontinuation of treatment or death (Dutcher et al., 2014). These toxicities are particularly apparent with the high doses of IL-2 that are required to stimulate CD8 T cell proliferation. A potential explanation for the complexities in outcome following IL-2 treatment is the lack of specificity, with an active area of research aiming to improve therapeutic IL-2 by altering its affinity for its receptors (Abbas et al., 2018; Boyman et al., 2006; Letourneau et al., 2010). Alternatively, contextual effects, arising from conditional dependence of signals, may explain the unexpected clinical effects. Sufficient understanding of the IL-2 homeostatic network is, however, first required to determine whether contextual signaling is involved.

The detailed study of the responsiveness to IL-2 of individual cell types has focused on exogenous provision to the desired therapeutic targets of Treg and CD8 T cells. A systematic network analysis of IL-2 sources and effects has been lacking. For successful utilization of IL-2 in the clinic, this immunological network understanding is critical. Here we developed a novel mouse strain for dissecting IL-2 network effects, and found that the biological effect of IL-2 differs markedly based on the cellular source. Even within the most studied axes of the IL-2 network, the responsiveness of Treg and CD8, new modalities were determined, with preferential responsiveness by Treg to IL-2 delivered in *trans* and by CD8 to IL-2 delivered in *cis*. Alternative contexts for IL-2 provision resulted in new IL-2-dependent biological circuits arising, the most notable being the dramatic expansion of eosinophils and CD8 Tregs following local IL-2 delivery by B cells. These results have profound implications for the clinical delivery of IL-2, with the potential to tailor immunological outcomes by altering the context of delivery rather than manipulating the molecular characteristics.

## Results

### A genetic switch for rewiring of the IL-2 production network

To define the sources of IL-2 production in the homeostatic system, we developed a highly sensitive flow cytometry protocol for detecting IL-2 production (**Figure S1**). As previously reported, conventional CD4 T cells demonstrated the highest potential for IL-2 production among the stimulated leukocyte lineages in the spleen,lymph nodes and lung tissue (**Figure 1A,B**). IL-2 production was also reliably detected in Foxp3^+^ Tregs, CD8 T cells, DN T cells, γδ T cells and innate lymphoid cells, expanding the potential sources of IL-2 to lineages previously thought to be silenced for this cytokine (**Figure 1A**). Based on the frequency of these lineages, conventional CD4 T cells comprised ~65% of all IL-2-producing cells in the spleen and lymph nodes, with CD8 T cells much of the remainder (**Figure 1B**). As this system depends on stimulated production, we also assessed sources of IL-2 production using an IL-2 fate-mapping system (*Il2^Cre^ Rosa^RFP^* mice) and *Il2^GFP^* reporter mice. Both systems confirmed the diversity of IL-2 sources, with, indeed, a higher fraction of IL-2 production coming from cells other than conventional CD4 T cells in these stimulation-independent assays (**Figure 1C,D, Figure S2**). Notably, the contribution of non-traditional cell types to IL-2 production was greater in non-lymphoid tissues such as the lung (**Figure 1A-D, Figure S2**). Corresponding mapping of IL-2 receptor expression identified Tregs as the dominant high-affinity receptor expressers, diverse lineages expressing the intermediate receptor (γδ T cells, CD8 T cells, DN T cells, NK cells, ILCs in secondary lymphoid tissues) and few lineages expressing the low affinity receptor, mainly DCs and lung ILCs (**Figure 1E**). In order to perturb this IL-2 production network, we developed a genetic switch for IL-2 expression. Using the constitutive *Rosa26* promoter and a floxed-stop expression system (**Figure 1F, Figure S3**), we created a system where Cre expression would induce cell lineage-specific IL-2 expression. The weak endogenous *Rosa26* promoter was used to ensure that IL-2 expression remained at physiological levels; indeed, transgenic IL-2 expression in this system was ~5% of the expression level of native IL-2 production by conventional CD4 T cells (**Figure 1G,H**). Together, these results identified a potentially complex IL-2 production network and the ability to perturb that network in a directed fashion.

**Figure 1.**
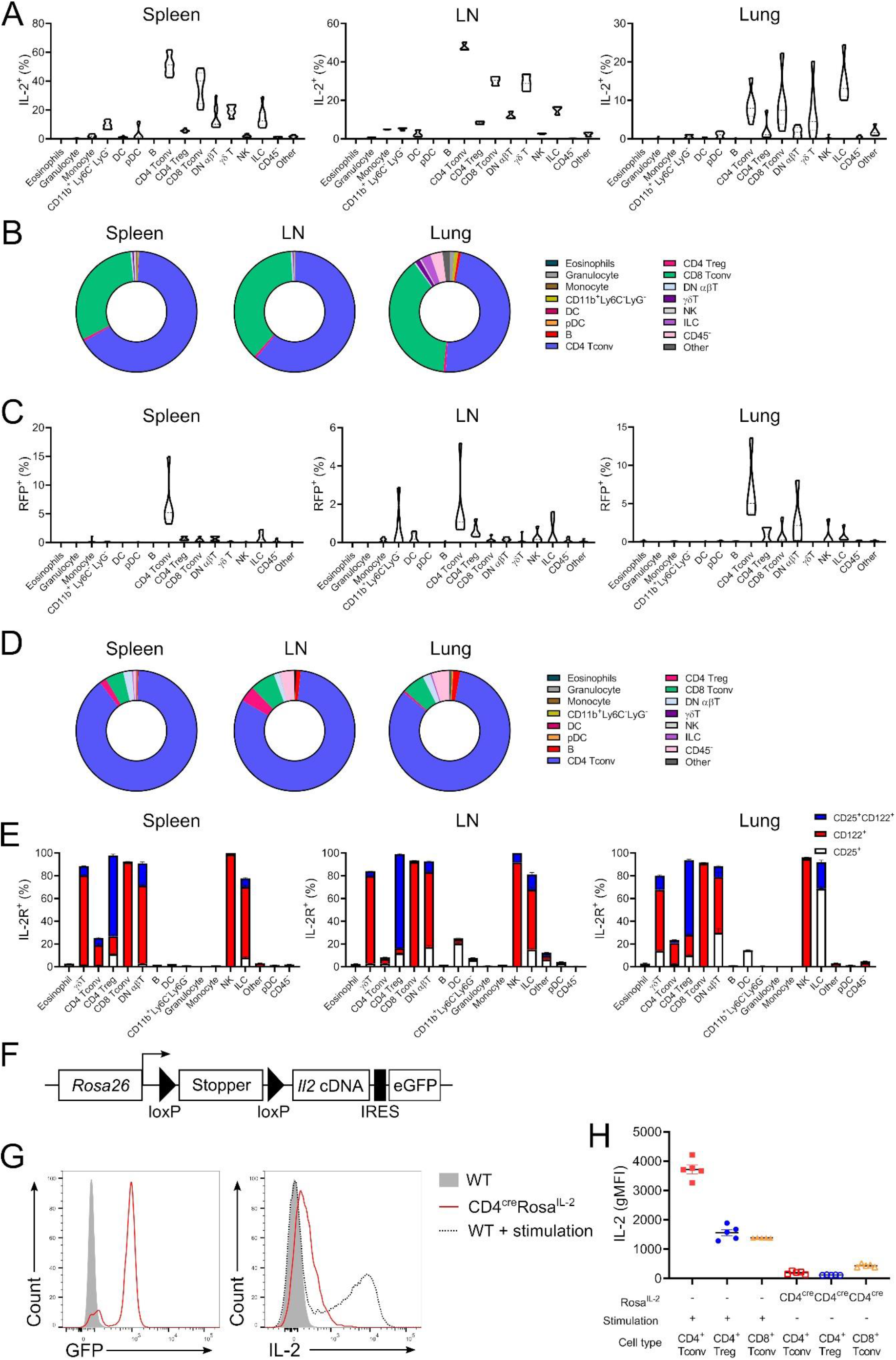
A complex production landscape for IL-2 in the homeostatic immune system. **(A)** IL-2 expression in spleen, LN and lung from WT cells stimulated *ex vivo* with PdBU/ionomycin for 4 h. n=6-9. **(B)** Frequency of each annotated cell type among the total IL-2^+^ population following *ex vivo* stimulation. **(C)** RFP expression in spleen, LN and lung from *Il2*^cre^*Rosa^RFP^* mice. n=6. **(D)** Frequency of each annotated cell type among the total RFP^+^ population from *Il2*^cre^*Rosa^RFP^* mice. **(E)** Expression of low-affinity (CD25^+^), intermediate-affinity (CD122^+^) or high-affinity (CD25^+^CD122^+^) IL-2 receptors in spleen, LN and lung WT cells. n=10. **(F)** Genetic construct of *Rosa^IL-2^* mice. **(G)** Representative histogram of GFP and IL-2 expression in CD4^+^ Tconv from wildtype (WT) and CD4^cre^Rosa^IL-2^ mice incubated for 4 h in the presence of Brefeldin A, or WT CD4^+^ Tconv stimulated for 4 h with PdbU/ionomycin/Brefeldin A. **(H)** gMFI of IL-2 expression from *ex vivo*-stimulated WT mice or non-stimulated CD4^cre^*Rosa^IL-2^* mice. n=5. Data are representative (A-B, G-H) or pooled (C-D) from at least two independent experiments.

### Inverted consequences of IL-2 production and response in CD4 and CD8 T cells drives differential network effects

The development of a Cre-inducible IL-2 system allowed us to constitutively drive IL-2 within the major IL-2-producing lineages. We first crossed the *Rosa^IL-2^* allele to CD4-Cre, active in both CD4 and CD8 T cells from the late double negative stage of thymic development, and the peripheral enhancer of CD8-Cre, active only in peripheral CD8 T cells. While the level of IL-2 produced by the genetic driver was much lower than the physiological capacity of these cells (**Figure 1G,H**), the system allows for constitutive expression, independent of antigen-mediated stimulation. Expression of constitutive low levelF IL-2 by CD8 T cells dramatically increased the cellularity of the spleen and lymph nodes (**Figure 2A**), largely through an expansion of the number of CD8 T cells and, to a lesser extent, Tregs (**Figure 2B,C**). Use of the CD4-Cre transgene surprisingly had a lower impact (**Figure 2A-C**), likely due to reduced thymopoiesis (**Figure S4**). Notably, both the relative and absolute number of conventional CD4 T cells collapsed following the provision of IL-2 either in *trans*, by CD8 T cells, or both in *cis* and *trans* (**Figure 2B,C**). This effect may have been mediated through an as yet undescribed toxic pathway of IL-2 on conventional CD4 T cells, or through the downstream effect of the large increase in Tregs, expanded to 70-90% of all CD4 T cells (**Figure 2D**). At a phenotypic level, CD4 and CD8 T cells were substantially altered by the IL-2 provision (**Figure 2E, Figure S5**). CD8 T cells suffered a relative loss of naïve T cells (**Figure 2F**), driven almost entirely by a substantial increase in IFNγ-producing central memory (CM) cells (**Figure 2F,G**). The collapse in CD4 T cell numbers was observed in both the CD4-Cre and CD8-Cre drivers. In the CD4-Cre driver only, this included a relative increase in activated CD4 T cells (**Figure 2H, Figure S5**), and an expansion of Th1 and Th17 cells (**Figure 2H,I**), although this was offset by the decrease in absolute number of CD4 T cells (**Figure 2C**). With both drivers, this expansion of IFNγ-producing CD8 TCM, of a magnitude akin to a lymphoproliferative disorder, was accompanied by fatality at around 4 months of age (**Figure 2J**), despite the increase in Treg numbers.

**Figure 2.**
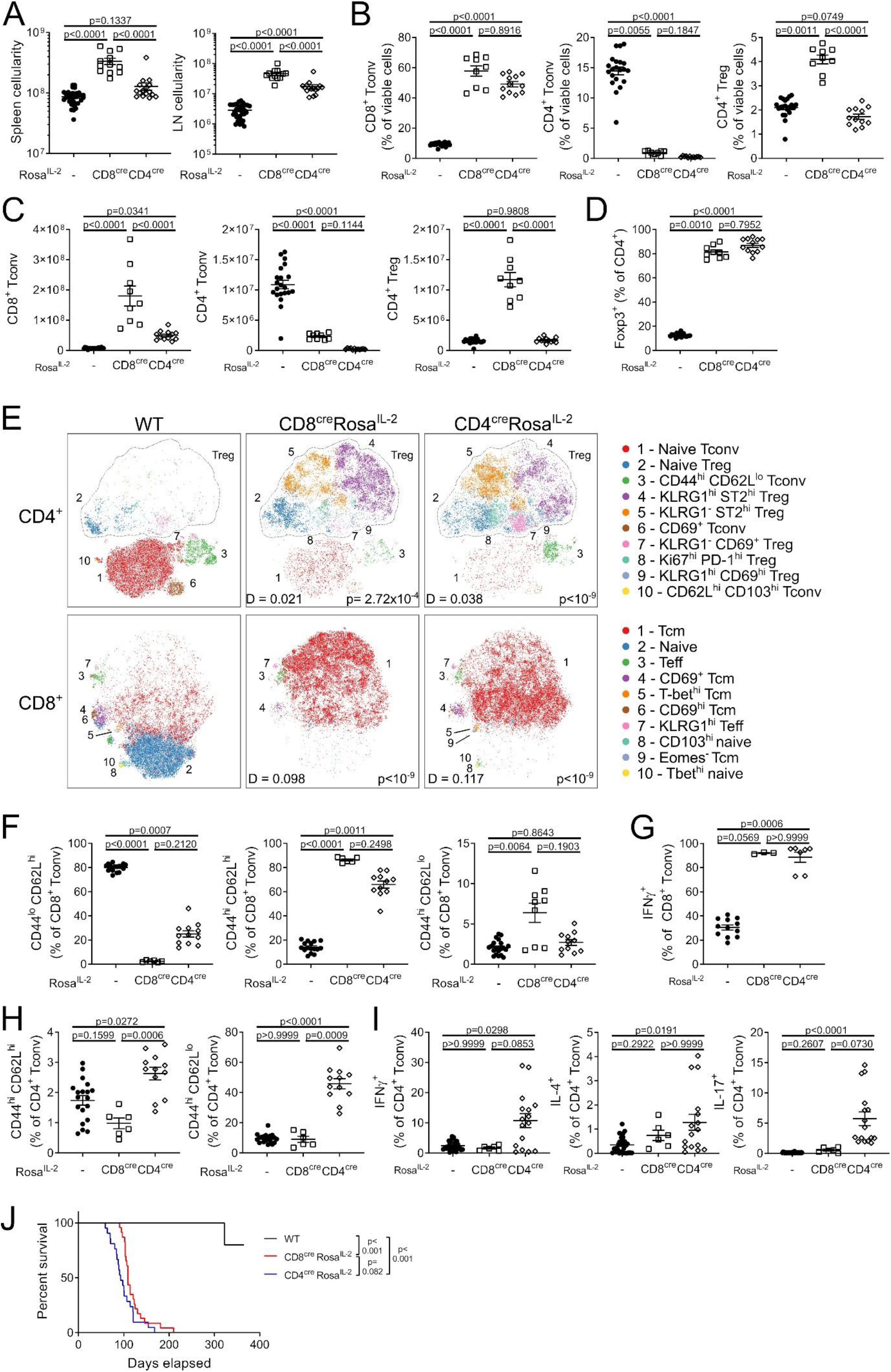
Fatal expansion of conventional CD8^+^ T cells follows IL-2 dysregulation despite heightened Treg representation. **(A)** Cellularity of spleen and LN of 4-6-week-old CD8^cre^ *Rosa^IL-2^*, CD4^cre^ Rosa^IL-2^ or littermate controls. n=12-34. **(B)** Frequency and **(C)** number of splenic CD8^+^ Tconv, CD4^+^ Tconv and CD4^+^ Treg. n=9-21. **(D)** Frequency of Foxp3^+^ cells among total CD4^+^ T cells. n=9-21. **(E**) tSNE representation of high-parameter flow cytometry data from splenic CD4^+^ (top) and CD8+ (bottom) T cells. FlowSOM clusters annotated based on differential expression of key markers. D value represents cross-entropy distance between samples. **(F)** Frequency of naïve, central memory (CD44^hi^ CD62L^hi^) or effector (CD44^hi^ CD62L^lo^) CD8^+^ Tconv. n=6-19. **(G)** IFNγ expression by CD8^+^ Tconv. n=3-12. **(H)** Frequency of central memory (CD44^hi^ CD62L^hi^) or effector (CD44^hi^ CD62L^lo^) CD4^+^ Tconv. n=6-19. **(I)** Cytokine production from CD4^+^ Tconv. n=6-32. **(J)** Survival analysis. n=5-23. Data representative (G) or pooled (A-F, H-J) from ≥ 2 independent experiments. Significance was tested by one-way ANOVA (A, C), Kruskal-Wallis (B, F-I), multiple Kolmogorov-Smirnov tests with Holm correction (E) or Mantel-Cox log-rank test (J).

The identification of a small population of IL-2-producing Tregs (**Figure 1A**) demonstrates that the reported *Il2* silencing through chromatin inaccessibility (Hemmers et al., 2019) is incomplete. While only ~1% of Tregs had a fate memory of IL-2 production in lymphoid organs, substantially higher numbers were observed in non-lymphoid environmental-interface tissues (**Figure 3A**). Phenotypic comparison of both IL-2 fate-mapped (**Figure 3B**) and stimulated IL-2 expressers (**Figure 3C**) demonstrated that the IL-2-producing Tregs were more likely to be activated and proliferating, suggesting loss of locus silencing is associated with stimulation. We therefore intercrossed the *Rosa^IL-2^* strain with *Foxp3^Cre^* mice (Rubtsov et al., 2008), to create a system where the silencing of *Il2* in Tregs was overridden (**Figure 3D**). A dose-dependent effect on splenic cellularity and mouse mortality (**Figure 3E,F**) was observed, with *Foxp3^Cre/wt^* female mice (in which only 50% of Tregs would activate *Rosa^IL-2^*. due to X chromosome inactivation) remaining healthy until nearly a year of age, while *Foxp3^Cre^* male mice (with 100% Treg activation of *Rosa^IL-2^*) developed lethal lymphoproliferation at ~5 months of age. Immunological assessment identified a largely stable leukocyte composition (**Figure 3G, Figure S6**). In line with the effect on mortality, it was only in the *Foxp3^Cre^ Rosa^IL-2^* mice that CD8 T cell numbers rose and CD4 T cell numbers collapsed (**Figure 3H**). Conversely, Treg numbers gave a graduated response in numbers (**Figure 3H,I**), while remaining phenotypically similar (**Figure 3J**). These system-wide data are consistent with prior work demonstrating priority access of Tregs to IL-2, and identify a threshold of cellular IL-2 provision at which CD8 T cells can become major IL-2-responders.

**Figure 3.**
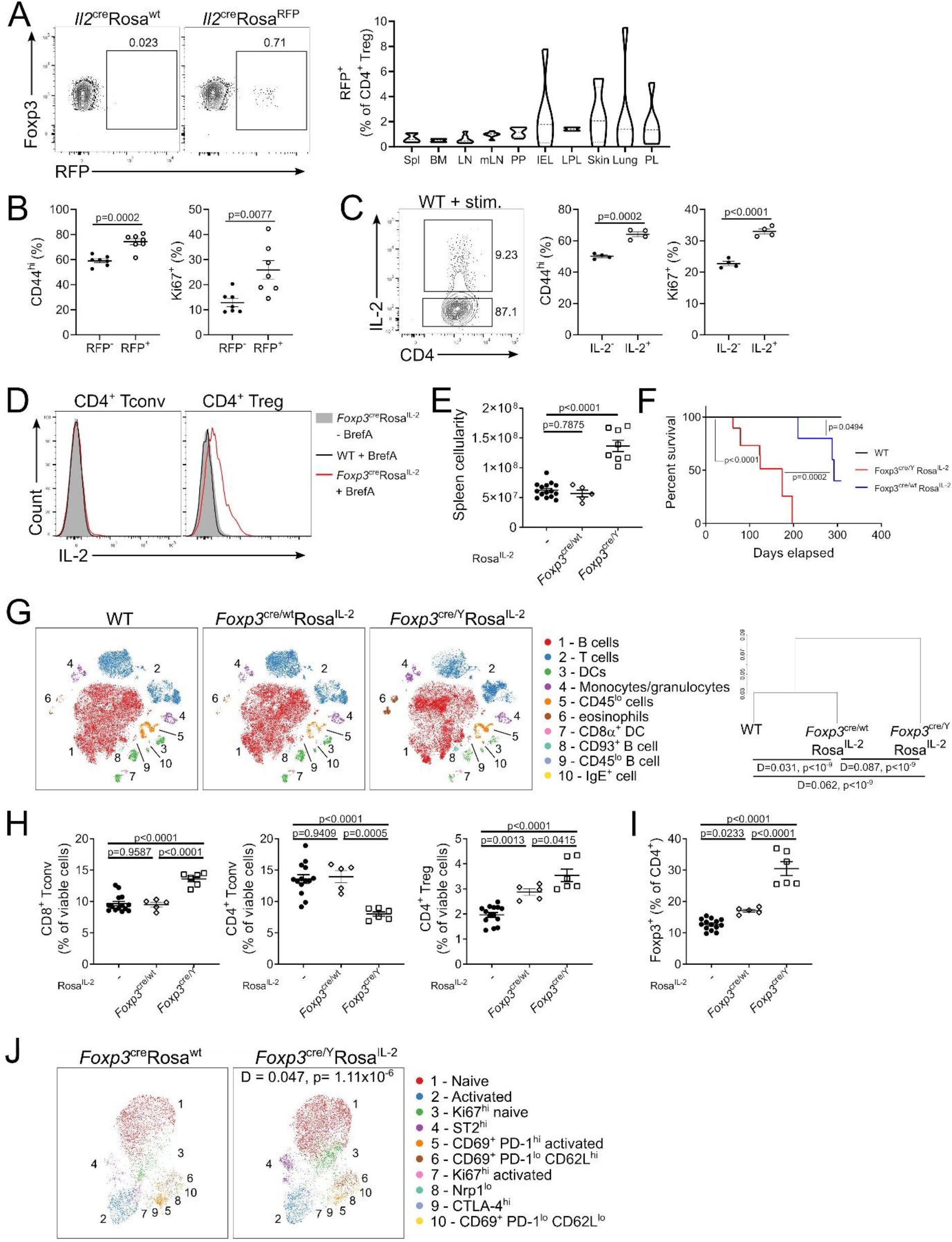
Treg self-sufficiency for IL-2 drives dose-dependent expansion. **(A)** IL-2 expression by CD4^+^ Treg from *Il2*^cre^Rosa^RFP^ mice. n=6. **(B)** Frequency of CD44 and Ki67 expression in Treg from *Il2*^cre^Rosa^RFP^ mice. n=7. **(C)** Frequency of CD44 and Ki67 expression in IL-2+/- Treg from *ex vivo* stimulation of WT cells. n=4. **(D)** Representative histograms of IL-2 expression in WT or *Foxp3*^cre^Rosa^IL-2^ mice incubated with Brefeldin A for 4 hours to prevent cytokine secretion. **(E)** Splenic cellularity of *Foxp3*^cre/wt^*Rosa^IL-2^*, *Foxp3*^cre^*Rosa^IL-2^* or littermate controls. n=5-15. **(F)** Survival analysis. n=11-23. **(G)** tSNE representation of high-parameter flow cytometry data from viable splenocytes (left). FlowSOM clusters annotated based on differential expression of key markers. Dendrogram showing comparative similarity calculated using cross-entropy distributions from tSNE (right). **(H)** Frequency of splenic CD8^+^ Tconv, CD4^+^ Tconv and CD4^+^ Treg. n=5-15. **(I)** Frequency of Foxp3^+^ cells among total CD4^+^ T cells. n=5-15. **(J)** UMAP representation of high-parameter flow cytometry data of *Foxp3*^cre^-expressing Treg from *Foxp3*^cre/wt^Rosa^wt^ or *Foxp3*^cre/wt^Rosa^IL-2^ mice. FlowSOM clusters annotated based on differential expression of key markers. D value represents cross-entropy distance between samples. Data are pooled (A-B, E-J) or representative (C-D) from at least two independent experiments. Significance was tested by paired t-test (B, C), one-way ANOVA (E, H-I), Mantel-Cox log-rank test (F), multiple Kolmogorov-Smirnov test with Holm correction (G) or two-sample Kolmogorov-Smirnov test (J).

The titrated response of Tregs to *cis*-production of IL-2 suggests that Tregs are more responsive to overall levels of IL-2 production than to autocrine sources. To directly test this hypothesis we created a titration of CD4-Cre *Rosa^IL-2^* bone-marrow with congenically-labelled wildtype cells (**Figure 4A**). The system allows the simultaneous dissection of dose-response effects and autocrine/paracrine effects. The dose-response effects demonstrated a linear-phase between 0-40% CD4-Cre *Rosa^IL-2^* bone-marrow chimerism, where additional IL-2-producing T cells resulted in increased Treg and decreased CD4 T cells, followed by a early plateau phase where additional IL-2 did not perturb the system further (**Figure 4B**). CD8 T cells also increased in number with higher representation of IL-2-producing bone-marrow (**Figure 4B**), suggesting a dose-dependent response for CD8 lymphoproliferation. The use of congenic bone-marrow to titrate allowed us to determine whether there was an additional benefit of autocrine IL-2 production (i.e., a disproportionate increase in cells originating from CD4-Cre *Rosa^IL-2^* bone-marrow) in addition to the dose-dependent effects. Compared to B cells as an internal control, CD8 T cells demonstrated a clear autocrine advantage when genetically licenced to produce IL-2 in an activation-independent manner (**Figure 4C**). Consistent with data from the CD8-Cre *Rosa^IL-2^* mouse, the TCM CD8 T cell subset had a preferential autocrine advantage (**Figure 4D, Figure S7**). CD4 T cells, by contrast, exhibited a clear autocrine disadvantage (**Figure 4C**), demonstrating that the reduction of CD4 T cells observed in IL-2-producing strains was not due to excessive suppression by Tregs, but rather due to a novel mechanism of autocrine IL-2-mediated negative feedback. Surprisingly, while Treg numbers expanded in response to more IL-2 production (**Figure 4B**), not only was no autocrine advantage observed, but a strong autocrine disadvantage was displayed (**Figure 4C**). This autocrine disadvantage was disproportionately observed among Tregs with a tissue-resident phenotype (**Figure 4E, Figure S7**). To identify the source of the competitive fitness disadvantage exhibited by IL-2-producing CD4 T cells, we investigated cytokine receptor expression in CD4-Cre *Rosa^IL-2^* mice. Compared to wildtype mice, CD25 was elevated in Treg and, to a lesser extent, conventional CD4 T cells in CD4-Cre *Rosa^IL-2^* mice (**Figure S8**). By contrast, expression of CD127 (IL-7Rα) was impaired on IL-2-producing conventional CD4 T cells but elevated on IL-2-producing CD8 T cells (**Figure 4F-H**). The net effect *in vivo* was a decrease in STAT3 Y705 and STAT5 Y694 phosphorylation in IL-2-producing conventional CD4 T cells, while cytokine signaling capacity was maintained or increased in IL-2 producing Tregs and CD8 T cells (**Figure 4I**). *In vitro* cytokine stimulation experiments indicated a pan-T cell defect in responding to IL-7 or IL-15 in IL-2-producing Treg, conventional CD4 T cells or CD8 T cells (**Figure 4J**). *In vivo*, we generated 50%:50% mixed bone-marrow chimeras of wildtype and CD4-Cre *Il2^flox^* mice, allowing the comparison of –IL-2-competent and IL-2-incompetent T cells in the same physiological environment. While IL-2 receptor expression was intact, the loss of IL-2 production allowed IL-7Rα expression to increase on conventional CD4 T cells (**Figure 4K**). Together these results suggest that IL-2 production by T cells comes at a cost of loss of sensitivity to IL-7. With the net impact of elevated IL-2 signaling and reduced IL-7 signaling having divergent outcomes in CD4 and CD8 T cells, autocrine IL-2 production preferentially expands CD8 T cells while contracting CD4 T cells.

**Figure 4.**
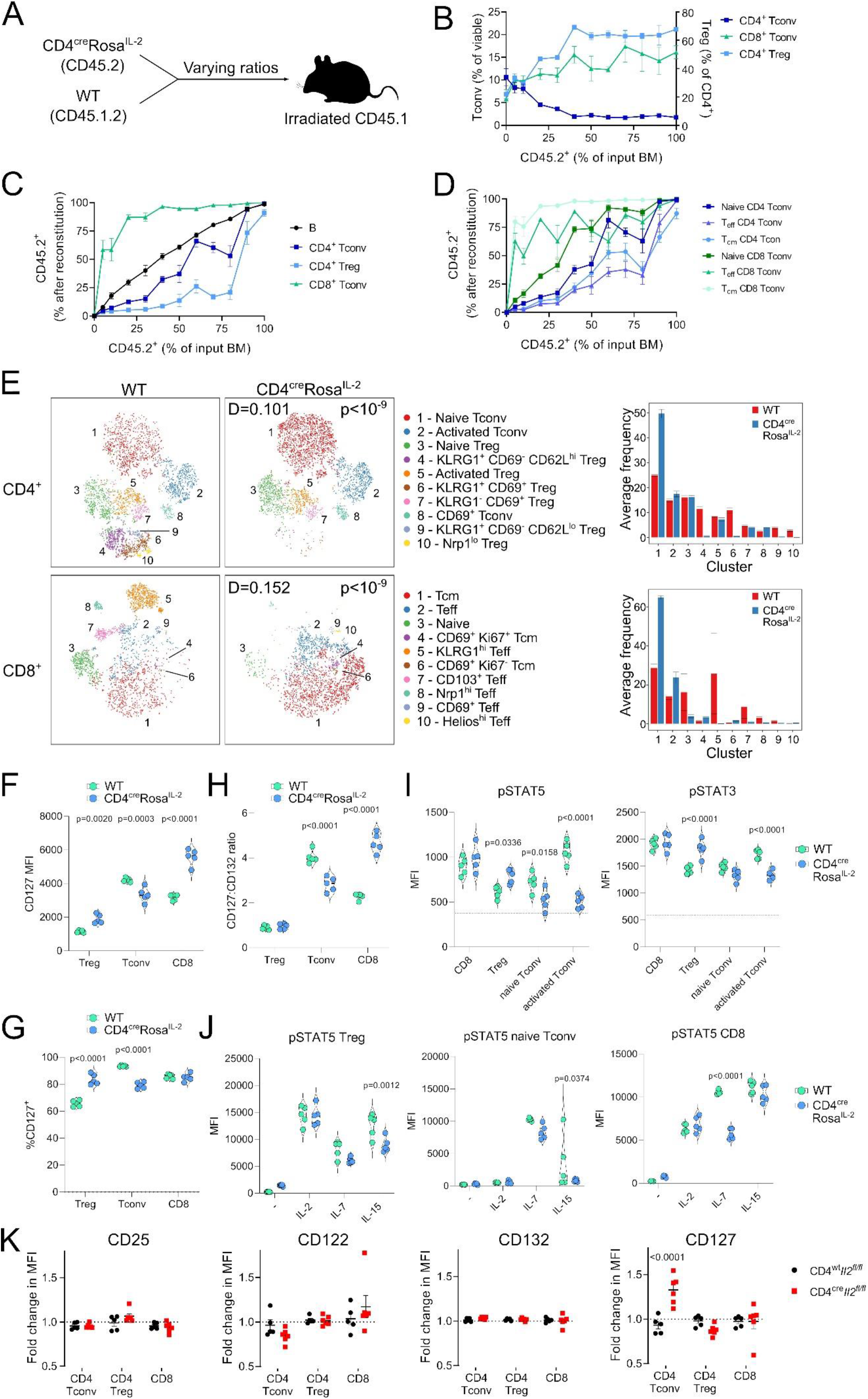
Autocrine IL-2 production drives divergent responses in CD4^+^ and CD8^+^ T cells. Chimeric mice were generated using bone marrow from WT (CD45.1^+^) and CD4^cre^*Rosa^IL-2^* (CD45.2^+^) at varying ratios. n=4-6, pooled from two independent experiments. **(A)** Schematic of experimental outline. **(B)** Frequency of CD4^+^ and CD8^+^ Tconv (as percentage of viable splenocytes) or CD4^+^ Treg (as percentage of total CD4^+^ T cells). **(C)** Frequency of splenocyte population derived from CD4^cre^*Rosa^IL-2^* (CD45.2^+^) bone marrow after reconstitution versus frequency of input bone marrow. **(D)** Frequency of naïve (CD44^lo^ CD62L^hi^), T_eff_ (CD44^hi^ CD62L^lo^) or T_cm_ (CD44^hi^ CD62L^hi^) derived from *CD4*^cre^*Rosa^IL-2^* (CD45.2^+^) in spleen. **(E)** tSNE representation of high-parameter flow cytometry data from splenic CD4^+^ (top) and CD8+ (bottom) T cells from 50% WT: 50% CD4^cre^*Rosa^IL-2^* chimeras. FlowSOM clusters annotated based on differential expression of key markers. D value represents cross-entropy distance between samples. **(F)** MFI and **(G)** percent of cells expressing CD127 (IL-7Rα) on CD4^+^ Treg, CD4^+^ Tconv and CD8^+^ T cells in WT and CD4^Cre^ Rosa^IL-2^ transgenic mice. (**H)** Ratio of CD127 MFI to CD132 (common γ chain) MFI. **(I)** pSTAT5 and pSTAT3 in freshly isolated T cells. FMO level indicated by dashed line. **(J)** Upregulation of pSTAT5 in response to cytokine stimulation in CD4^+^ Treg, naïve CD4^+^ Tconv and CD8^+^ T cells. **(K)** Fold change in MFI of receptor expression in CD45.2^+^ cells to CD45.1^+^ cells in chimeric mice generated as 50% CD4^cr^*Il2^fl/lf^* or CD4^wt^*Il2^fl/fl^* (CD45.2^+^):50% WT (CD45.1^+^). Significance was tested by two-sample Kolmogorov-Smirnov test (E), Sidak’s multiple comparison test on 2-way ANOVA (F-K).

### Context-dependent perturbation of IL-2 production reveals novel cellular circuits

To test for contextual symmetry in the IL-2 network, we amplified three existing but minor IL-2 sources: dendritic cells (DCs), NK cells and B cells, through crossing the *Rosa^IL-2^* allele onto the *Clec9a^Cre^, Ncr1^Cre^* (NKp46-Cre) and *Cd19^Cre^* strains, respectively. *Clec9a^Cre^ Rosa^IL-2^* mice, with basal production of IL-2 from DCs, maintained a normal number of DCs, with the exception of a large increase in the CD103^+^ population (**Figure S9**). Responding populations were largely restricted to Tregs, in particular those with a resident-like phenotype (**Figure S9**). Unlike the observations using T cell-drivers of IL-2, CD8 T cells were reduced rather than expanded, suggesting a primary wiring of DC-derived IL-2 to the Treg sink, consistent with the phenotype of CD11c-Cre IL-2^flox^ mice (Owen et al., 2018). NKp46-Cre *Rosa^IL-2^* mice, with basal production of IL-2 from NK cells and ILC3s demonstrated a large (~10-fold) and specific expansion of the NK population, with only a minor increase in Treg numbers and a small decline in conventional CD4 and CD8 numbers (**Figure S10**). These data suggest that, like CD8 T cells, NK cells preferentially respond to intrinsic production of IL-2.

Surprisingly, production of IL-2 from B cells, using *Cd19^Cre^ Rosa^IL-2^* mice, drove a distinct cellular network to all other IL-2 sources, despite similar levels of net IL-2 production to CD4^Cre^ *Rosa^IL-2^* mice (**Figure 5A**). *Cd19^Cre^ Rosa^IL-2^* mice exhibited an enlarged spleen, but with reduced splenic cellularity due to extensive fibrosis (**Figure 5B**). Assessment of leukocyte changes identified the primary shift as a 50-fold increase in eosinophils (**Figure 5C,D**). Eosinophilia was accompanied by large increases in IL-13 and, especially, IL-5 (**Figure 5E**). ILC were the dominant source of IL-5 in *Cd19^Cre^ Rosa^IL-2^* mice (**Figure 5F**), with ILC2 numbers expanded 100-fold, to compromise >90% of the ILC pool (**Figure 5G,H**). A similar increase in ILC2 and eosinophils was observed in the bone-marrow (**Figure S11**). To test the possibility that this phenotype is driven by early bone-marrow expression of IL-2, rather than B cell-specific expression, we made the additional crosses to *CD23^Cre^* and *Osx^Cre^* mice. *CD23^Cre^*, despite turning on later in mature B cells, generated the same eosinophil-dominated phenotype (**Figure S12**), while *Osx^Cre^*, active in bone-marrow osteoblasts, did not (**Figure S13**). Neutralization of IL-5 in *Cd19^Cre^ Rosa^IL-2^*mice prevented the eosinophilia (**Figure 5I**), demonstrating that B cell-specific production of IL-2 initiated an unconventional cellular circuit expanding IL-5-expressing ILC2s and, downstream, eosinophils. This immune dysregulation did not result in significant excess mortality (**Figure 5K**), unlike that observed with the T cell Cre-drivers (**Figure 2J, Figure 3F**).

**Figure 5.**
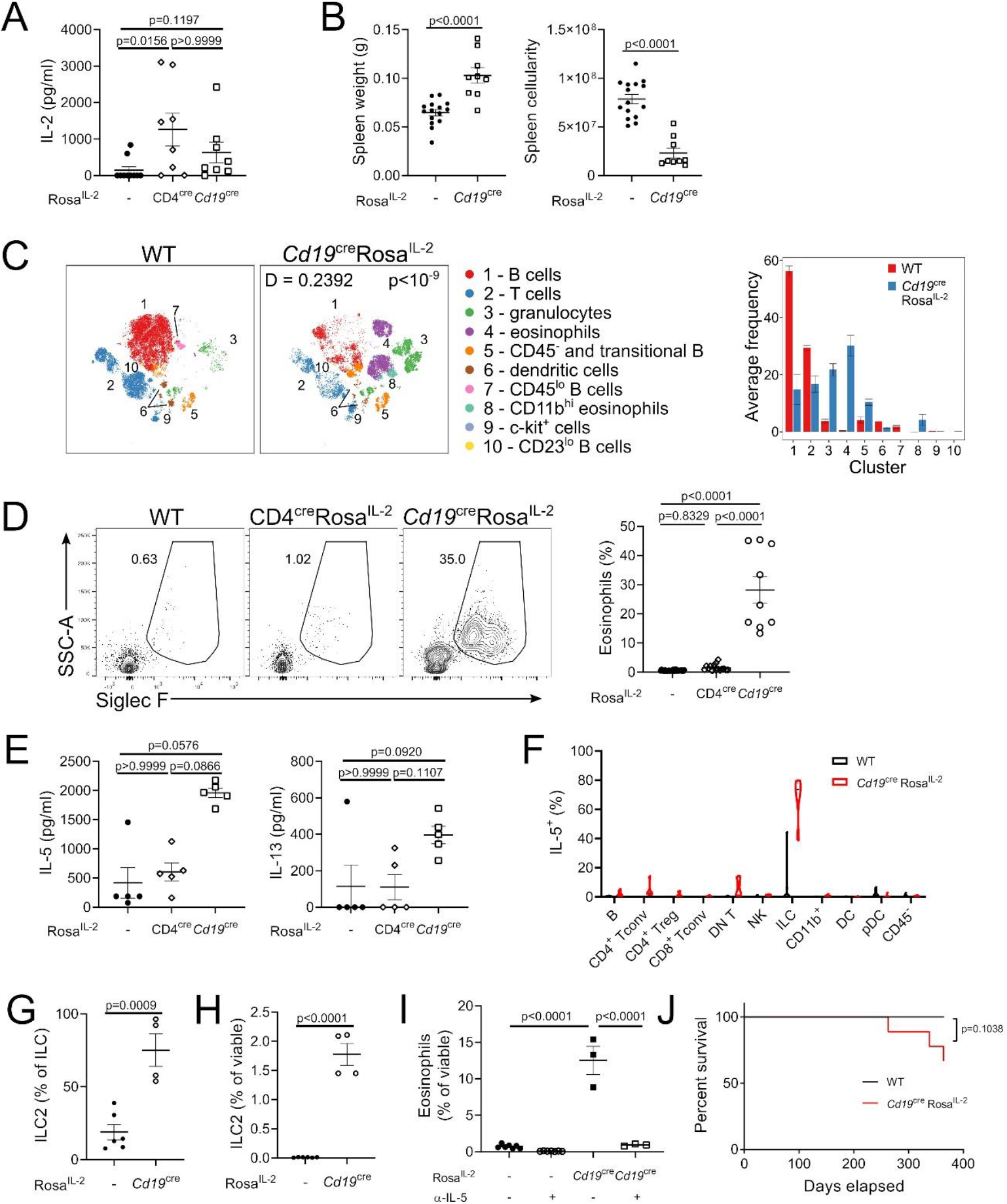
Expression of IL-2 by B cells drives a distinct ILC2-eosinophil-orientated cellular circuit. **(A)** Luminex analysis of IL-2 cytokine in serum. n=8-10. **(B)** Spleen weight and cellularity of 4-6-week-old *Cd19*^cre^ *Rosa^IL-2^* and littermate controls. n=9-15. **(C)** Representative tSNE of high-parameter flow cytometry data from splenic viable cells and average frequency of each cluster per mouse. **(D)** Representative gating and frequency of eosinophils in spleen. n=9-29. **(E)** Luminex analysis of indicated cytokine in serum. n=5. **(F)** IL-5 expression among cell lineages in spleen. n=6-9. **(G)** Frequency of GATA3^+^ ILC2 among total ILCs in spleen. n=4-6. **(H)** Frequency of ILC2 in spleen. n=4-6. **(I)** Frequency of splenic eosinophils in mice treated with anti-IL-5 neutralising antibody. n=3-8. **(J)** Survival analysis. n=7-9. Data pooled (A-D, F-I) or representative (E) from at least 2 independent experiments. Significance was tested by one-way ANOVA (A, D, E, I), unpaired t-test (B, G, H), two-sample Kolmogorov-Smirnov test (C) or Mantel-Cox log-rank test (J).

Finally, we observed a second major abnormality in *Cd19^Cre^ Rosa^IL-2^* mice: a 40-fold expansion of CD8^+^ Foxp3^+^ cells (**Figure 6A,B**). This population, exceedingly rare in wildtype mice, expanded to ~20% of CD8 T cells in *Cd19^Cre^ Rosa^IL-2^* mice. Foxp3 expression in CD8^+^ Foxp3^+^ cells correlated with three independent Foxp3 marker systems (**Figure S14**). Notably, the provision of IL-2 to drive this increase needed to come from B cells, as the population remained small in mice with the CD4-Cre or CD8-Cre drivers (**Figure 6B,C**). CD8^+^ Foxp3^+^ cells displayed a distinct phenotype from either conventional CD8 T cells or CD4 Tregs (**Figure 6D, Figure S14**), with expression of some Treg markers, such as CD25, but low expression of others, such as Nrp1 (**Figure 6E**). The CD8^+^ Foxp3^+^ cells expanded in *Cd19^Cre^ Rosa^IL-2^* mice were, however, *bona fide* Tregs, with an *in vitro* suppressive capacity on both CD4 and CD8 conventional T cells indistinguishable from that of CD4^+^ Foxp3^+^ cells (**Figure 6F**). Analysis of the small CD8^+^ Foxp3^+^ population present in wildtype mice demonstrated that they were CD8αβ cells, however, and unusually, the CD8-Cre transgene had poor penetrance in the CD8^+^ Foxp3^+^ population (**Figure S14**). As this was akin to CD4^+^CD8^+^ double positive thymocytes, we sought to determine whether this population were CD8^+^ MHCII-restricted cells. Comparison of wildtype, CD1d KO, MHCI KO, MHCII KO and MHC I/II double KO mice demonstrated that CD8^+^ Foxp3^+^ are classical MHCI-restricted CD8 T cells (**Figure 6G, Figure S15**). High dimensional flow cytometry profiling of CD8^+^ Foxp3^+^ cells in wildtype mice found a highly skewed subset distribution compared to conventional CD8^+^ Foxp3^−^ cells (**Figure 6H**). In particular, splenic CD8^+^ Foxp3^+^ cells were greatly enriched (~50%) for the CXCR5^+^PD1^+^ subset (**Figure 6I,J**). As this phenotype is shared with follicular helper CD4 T cells (Vinuesa et al., 2016), we investigated the anatomical distribution of CD8^+^ Foxp3^+^ cells in the spleen, and found that while <1% of CD8 T cells in total, this population constituted ~60% of CD8 T cells present in the B cell zone of naïve mice (**Figure 6K**). These data provide a plausible route for the specific expansion of CD8 Tregs in *Cd19^Cre^ Rosa^IL-2^* mice: their presence in the B cell zone puts them in close proximity with B cell-derived IL-2 production, while making them relatively refractory to T cell-derived IL-2. These mice, in addition to providing for sizeable populations of CD8 Tregs amenable to functional assays, therefore illustrate the contextual importance of IL-2 production.

**Figure 6.**
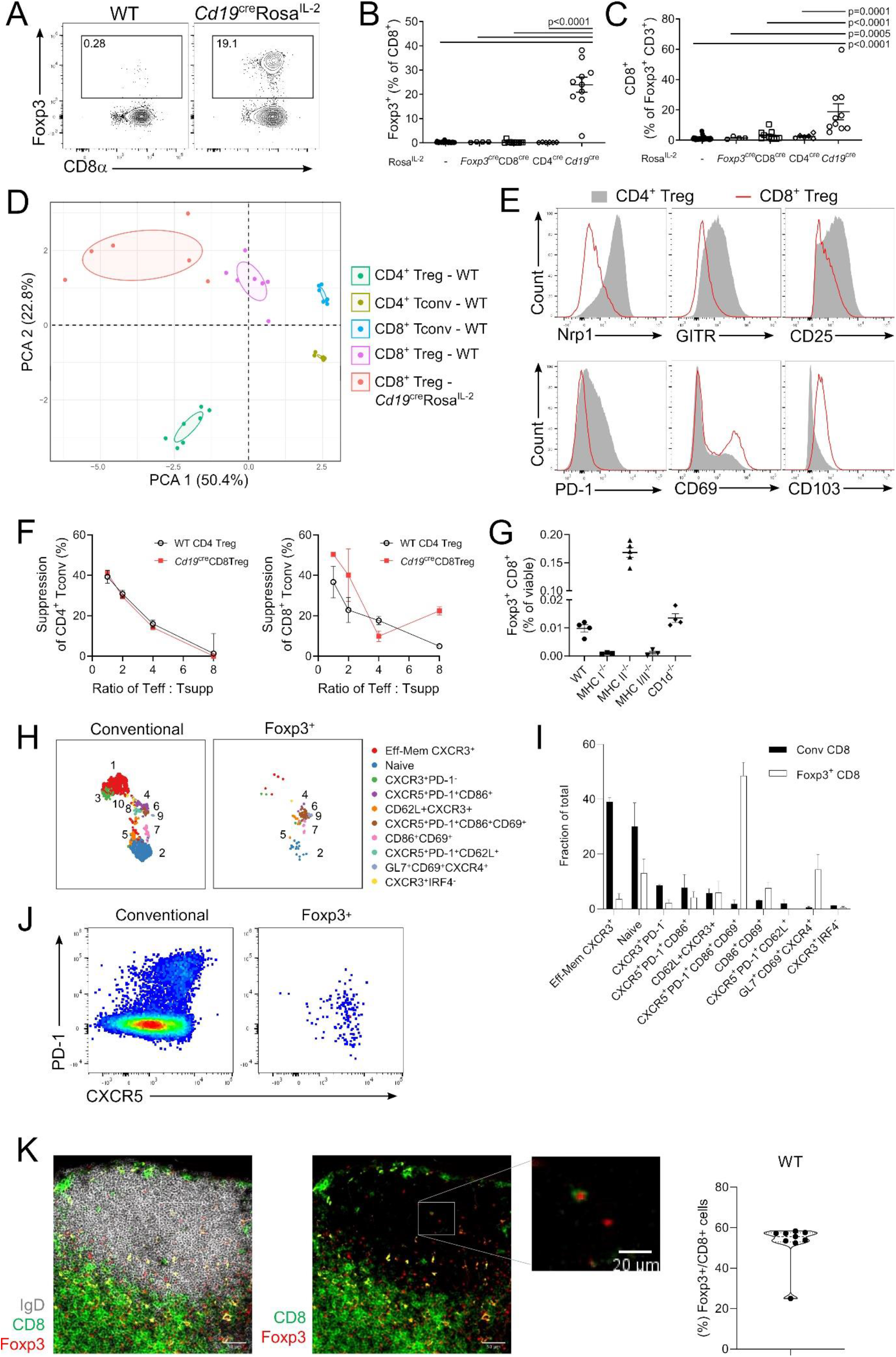
CD8^+^Foxp3^+^ Tregs are revealed through B cell production of IL-2. **(A)** Representative gating of Foxp3^+^ CD8^+^ T cells in *Cd19*^cre^Rosa^IL-2^ mice and littermate controls. **(B)** Frequency of Foxp3 expression among CD8^+^ T cells in Rosa^IL-2^ strains. n=6-32. **(C)** Frequency of CD8^+^ cells among total Foxp3^+^ T cells. n=9-32. **(D)** Principal component analysis of flow cytometric markers on splenic T cell populations. **(E)** Representative expression of indicated markers on CD4^+^ Treg or CD8^+^ Treg from *Cd19*^cre^Rosa^IL-2^ mice. **(F)** *In vitro* suppression assay comparing ability of CD4^+^ Treg and CD8^+^ Treg to suppress CD4^+^ Tconv (left) or CD8^+^ T conv (right) proliferation. **(G)** Frequency of Foxp3^+^ CD8^+^ cells in spleens of wildtype, *MHCI^-/-^, MHCII^-/-^, MHCI^-/-^MHCII^-/-^* and *CD1d^-/-^* mice. **(H)** UMAP of high-parameter flow cytometry data from conventional (Foxp3^−^) CD8^+^ T cells and CD8^+^ Tregs, with **(I)** frequency distribution of Foxp3^+^ and conventional CD8 across FlowSOM clusters (n=3). **(J)** Representative flow plot showing PD-1 and CXCR5 expression. **(K)** Representative immunofluorescence staining of CD8α, Foxp3 and IgD of wildtype lymph nodes (scale bars: 50μm, 20μm for inset), with quantification of Foxp3^+^ cells among total CD8 T cells in the B follicle (n=9). Data pooled (B, C, D, G, H, I) or representative (A, E, F, J, K) from at least two independent experiments. Significance was tested by one-way ANOVA (B, C, G).

## Discussion

The IL-2 network includes a diverse array of cellular sources and multiple potential responder cell types. Under a standard “immunology as a single cell suspension” perspective, IL-2 generated by any source enters a common pool, with consumption driven by affinity-based cellular capture. The dominant components of the IL-2 network at homeostasis are compatible with this simplified model. Conventional CD4 T cells are the dominant source, and the two main sinks are Tregs and CD8 T cells. Priority within the sink populations is consistent with the affinity of receptor expression, with Tregs responding at a lower dose (and expressing the high affinity trimeric receptor) and CD8 T cells responding at higher doses (consistent with expression of the medium affinity dimeric receptor). Amplification of minor sources of IL-2, however, demonstrates the flaws inherent to the context-independent model. NK cells, ILC2s (Spolski et al., 2018) and CD8 Tregs (current study) can all express the high affinity trimeric receptor, and yet do not respond to T cell-derived IL-2. Instead, NK cells responded only to autocrine production while ILC2s and CD8 Tregs were expanded only by B cell-sourced IL-2. This demonstrates that the range of responses possible to IL-2 is not merely due to receptor expression and preferential capture based on affinity, but rather there is a context-dependent component. While we do not negate the utility of the affinity-based competition model, this data does necessitate the inclusion of context-sensitivity into the model, where the cellular source of IL-2 dictates its function. In the absence of evidence for more exotic hypotheses, such as post-transcriptional modification by the cellular source and binding to undiscovered alternative receptors (as commonly occurs for chemokines, altering receptor preference (Stone et al., 2017)), the parsimonious explanation lies in a proximity-based effect. The diffusion radius of IL-2 is as low as 30μm with high densities of consuming cells (Oyler-Yaniv et al., 2017), which would be expected to result in a sharp IL-2 gradient at the border of the T cell zone. The expansion of ILC2s and CD8 Tregs supports anatomical proximity as the contextual source, as we find CD8 Tregs heavily enriched in the B cell zone, while ILC2 reside in the B cell-adjacent interfollicular area (Mackley et al., 2015), consistent with their function in promoting early antibody responses (Drake et al., 2016). Proximity-based contextualism would likely result in different IL-2 circuitry in non-lymphoid organs, such as the ILC3 to Treg link proposed in the gut (Zhou et al., 2019). IL-2 may itself contribute to re-wiring of the circuit in such contexts, as IL-2 response genes are highly enriched for microenvironmental sensors (Rollings et al., 2018; Ross et al., 2016).

The relationship connecting CD4 T cell IL-2 production to Tregs and CD8 T cells is the best-described aspect of the IL-2 network. In the thymus, production of IL-2, primarily from DCs (Owen et al., 2018) and self-reactive thymocytes (Hemmers et al., 2019; Owen et al., 2018), helps drive Treg differentiation via signaling to the CNS2 Foxp3 genetic element (Feng et al., 2014). Intriguingly, a potential role for autocrine IL-2 in the early thymic Treg precursor has been identified (Chawla et al., 2020). In the periphery, IL-2 production from activated CD4 T cells is critical to support the fitness (Fontenot et al., 2005), survival (Pierson et al., 2013; Shi et al., 2018) and regulatory function (Chinen et al., 2016) of Tregs. CD8 T cells are highly dependent on IL-2 for the formation of memory (Williams et al., 2006), setting up a competitive dynamic between Tregs and CD8 T cells for IL-2 consumption (Pandiyan et al., 2007). Indeed, the preferential ability of Treg to capture IL-2, based on expression of CD25, impairs the ability of CD8 T cells to enter the memory fate (Chinen et al., 2016). Treg-based competition is, however, also beneficial to the quality of the CD8 T cell response, with more complete suppression of low affinity clones (Pace et al., 2012). The extraordinary capacity of both Tregs and CD8 T cells to respond to IL-2 dictates the highly-dependent nature these cells have on conventional CD4 T cells, their primary source. Such a system provides a failsafe to prevent runaway proliferation, as observed when we permit the normally forbidden constitutive expression in these restricted lineages. Notably, however, the outsourcing of IL-2 production to conventional CD4 T cells shifts the risk of a positive feedback loop onto the source cell type. Here the toxicity cost of IL-2 production, rather than being an inadvertent metabolic cost, may serve as an engineered regulatory check on a potential positive feedback loop. We identified *in vivo* the same effect previously observed *in vitro* of IL-2-mediated downregulation of IL-7 receptor expression (Xue et al., 2002). The ability of IL-2 to signal within the endosomal compartment (Konstantinidis et al., 2005; Volko et al., 2019) may provide the mechanistic route through which IL-2 production drives competitive costs in CD4 T cells, as the IL-7 receptor sequestration process could occur during the process of IL-2 secretion. This coupling of IL-2 production to IL-7 desensitization may thus provide a production-cost mechanism to IL-2-producing CD4 T cells. Intriguingly, the same endosomal signaling pathway could account for the competitive advantage when initiated in CD8 T cells, as the endosomal compartment concentrations may compensate for the lower affinity of the dimeric receptor. The network analysis thus reveals elegant details even for the best-described IL-2 cellular circuits.

Foxp3^+^ CD8 Tregs, distinct from other CD8 populations with proposed suppressive capacity, have been previously described in mouse and human (Mayer et al., 2011; Vieyra-Lobato et al., 2018). The CD8^+^Foxp3^+^ population is extremely low, 0.1% of CD8 T cells in mice and 0.3% in humans (Churlaud et al., 2015), although it has been reported to expand in response to IL-2 treatment (Rosenzwajg et al., 2015), allogeneic transplantation (Beres et al., 2012; Robb et al., 2012) and retrovirus infection (Nigam et al., 2010). The extremely low numbers have precluded ready functional validation or wide acceptance among the immunological community. The existence of a link between CD8 Tregs and B cells, identified here, provides both a novel insight into the biology of this neglected population and also a tool for study, allowing CD8 Tregs to be purified in numbers akin to CD4 Tregs.

The anomalous circuit driven by B cell-derived IL-2 production provides a mechanistic explanation for a long-standing enigma of IL-2 clinical use. In the first clinical trials, in the 1980s, of recombinant human IL-2 in primary immunodeficiency (Dopfer et al., 1984), AIDS (Kern et al., 1985) and cancer (Macdonald et al., 1990b), patients almost invariably developed eosinophilia, accompanied by high IL-5 titers. Despite treatment modification to reduce toxicity, similar results are a long-standing feature of IL-2 treatment for cancer, including pediatric tumors (Roper et al., 1992), renal cell carcinoma (Clark et al., 1999; Lee et al., 2010; Moroni et al., 2000), non-small cell lung cancer (Ardizzoni et al., 1994), neuroblastoma (Pardo et al., 1996), mesothelioma (Nano et al., 1998) and melanoma (Cragun et al., 2005; Woodson et al., 2004), with eosinophilia in treated patients being predictive of treatment failure (Moroni et al., 2000). Human eosinophils were reported to express IL-2 receptors (Rand et al., 1991), however *in vitro* assays suggested eosinophilia was due to IL-5, rather than direct effects of IL-2 (Macdonald et al., 1990a). IL-5-producing T cells were initially proposed as an intermediary (Enokihara et al., 1988; Enokihara et al., 1989), prior to the discovery of IL-5-producing ILC2s. Using a mouse model of IL-2-anti-IL-2 antibody complex injection, the Bluestone group demonstrated that ILC2s were the primary IL-5-expressing cell arising following these injections, and that ablation of all IL-5^+^ cells prevented eosinophilia (Van Gool et al., 2014). Our results explain why endogenous IL-2 production does not drive the same eosinophilic outcome, with only B cell-sourced IL-2 precipitating the ILC2 circuit. We propose that exogenous IL-2 provision violates the default anatomical restriction of major IL-2 production to T cell zones, thus precipitating a normally quarantined reaction. Intriguingly, one of the physiological contexts in which this B cell-driven circuit is naturally amplified may be that of parasitic infection. During infection with *Heligmosomoides polygyrus*, B cell production of IL-2 is required for parasitic control (Wojciechowski et al., 2009). While attributed to direct support for Th2 cells, IL-2 in this context may provide indirect support via ILC2s (Pelly et al., 2016). Eosinophilia in the context of *H. polygyrus* is superfluous for clearance (Urban et al., 1991), however amplification of this circuit would be beneficial in other helminthic infections (Klion and Nutman, 2004), demonstrating the contingent value of the B cell-ILC2-eosinophil circuit identified here.

A shift from an affinity-based competition model to a context-sensitive model creates new potentials for therapeutic delivery. The discrepancy between objective and outcome in IL-2 trials has been attributed to wide-spread receptor expression, with extensive research going into the design of synthetic IL-2 mutants (Spangler et al., 2015). The concept behind this approach is that by restricting IL-2 impact to only one receptor, adverse effects will be eliminated. IL-2 engineering is primarily achieved via altering binding to either CD25 or CD122. Reduced binding to CD25 (Carmenate et al., 2013) or enhanced binding to CD122 (Levin et al., 2012) promotes CD8 T cell responses, while reduced binding to CD122 (Peterson et al., 2018) or enhanced binding to CD25 (Rao et al., 2005; Rao et al., 2003) accentuates the Treg response. More elaborate engineering includes combining mutations (Sun et al., 2019), modifications to the common CD132 chain (Mitra et al., 2015), fusion to antibody domains (Khoryati et al., 2020; Spangler et al., 2018; Sun et al., 2019) or even *de novo* mimics that trigger receptor binding without homology to IL-2 (Silva et al., 2019). In each case, while the engineered properties impart receptor-specificity, they neglect contextual factors, and therefore cannot impart cellular-specificity. It is only when coupled with cell therapy that biochemical engineering of IL-2 can impart cell-specificity, such as through the generation of orthogonal IL-2 treatment coupled to transfer of cells engineered to express an orthogonal receptor (Sockolosky et al., 2018). An understanding of the contextual sensitivity of IL-2 may guide improved therapeutic bioengineering. For example, the initiation of the adverse eosinophilic circuit may be avoided by engineering exogenous IL-2 to recapitulate the quarantining of the B cell zone observed by the primary endogenous sources. Antibody-mediated guidance to T cell zones may achieve this goal, with therapies such as Darleukin (L19-IL-2 fusion) providing proof-of-concept for antibody-guided IL-2 delivery. Alternatively, the competitive advantage of CD8 T cells with self-production of IL-2, even within environments of enriched IL-2 available, suggest that targeting production to CD8 T cells themselves may drive the desired response, for instance in a tumor setting. Here approaches such as *in vivo* delivery of IL-2 plasmids (Lohr et al., 2001) could be coupled to CD8 T cell-specific promoters. A synthesis of biochemical engineering and context-sensitive design may unlock the long-awaited therapeutic potential of this key immunological player.

## Acknowledgements

This work was supported by the VIB, FWO (to A.L.), the ERC Consolidator Grant TissueTreg (to A.L.), Alzheimer’s Association Research Grant (to A.L.), and the Biotechnology and Biological Sciences Research Council (BBSRC) through Institute Strategic Program Grant funding BBS/E/B/000C0427 and BBS/E/B/000C0428, and the BBSRC Core Capability Grant to the Babraham Institute. K.S. was supported by a fellowship from Vetenskapsrådet, M.A. was supported by the French MS society (ARSEP). The authors acknowledge David Posner (Babraham Institute), Aleksandra Brajic, Qin Wu and Katinka van Dongen (VIB) for technical assistance, Jeason Haughton (VIB) and the BSU staff for animal curation, Rachael Walker and the Babraham Institute Flow Cytometry Core, Pier-Andrée Penttila and the KUL FACS Core, Babraham Institute Imaging Core, Sonia Agüera for scientific illustrations, Caetano Reis e Sousa (Francis Crick Institute), Michael Farrer (University of Minnesota), Carla Shatz (Stanford University) and Michelle Linterman, Rahul Roychoudhuri and Martin Turner (Babraham Institute) for the provision of mice and advice on experimental design.

## Author Contributions

Conceptualization: J.D. and A.L.; Methodology: C.E.W., K.S., O.T.B., M.A., A.M., L.K., T.Y.F.H., S.L., J.D. and A.L; Software: C.P.R.; Formal Analysis: C.E.W., O.T.B., C.P.R.; Investigation: C.E.W., O.T.B., K.S., F.L. and J.D.; Resources: T.H. and S.S.; Writing – Original Draft: C.E.W. and A.L.; Writing – Review & Editing: C.E.W, K.S., O.T.B., J.D. and A.L.; Visualization: C.E.W., C.P.R., J.D. and A.L.; Supervision: S.L., J.D. and A.L.; Project Administration and Funding Acquisition: A.L.

