## Supplementary methods and figures for "Context-dependent effects of IL-2 rewire immunity into distinct cellular circuits"

### *Mice*

The *Rosa*$^{IL\text{-}2}$ allele was generated by inserting the open reading frame of mouse *Il2* (transcript ID ENSMUST00000029275) into the first intron of the Rosa26 locus in C57BL/6N embryonic stem (ES) cells (Schoonjans et al., 2003). The gene targeting strategy is depicted in **Supplementary Figure 3**. The targeting construct consisted, from 5' to 3', of (i) 1082bp homologous region, (ii) adenovirus major late transcript splice acceptor (Friedrich and Soriano, 1991), (iii) LoxP-flanked neomycin resistance cassette (PGK promoter – NeoR/KanR – PGK polyadenylation signal), (iv) 3x SV40 polyadenylation signal, kozak-preceded *Il2* open reading frame, (v) IRES sequence, (vi) EGFP sequence, (vii) bovine growth hormone polyadenylation signal, (viii) 4264bp homologous region, and finally (ix) the diphtheria toxin subunit A gene (PGK promoter – DTA including SV40 small t antigen intron - bovine growth hormone polyadenylation signal) to select against random integration events. The construct was linearized with PvuI. The complete nucleotide sequence of the final targeting vector can be obtained from the authors upon request. Correctly targeted ES cell clones were identified by PCR with primer 5'- TAG GTA GGG GAT CGG GAC TCT-3' and 5'- GCG AAG AGT TTG TCC TCA ACC-3', site-specific integration was confirmed by Southern blotting and ES cells were injected into C57BL/6J albino blastocysts for the generation of chimeric mice. Chimeric males were crossed to C57Bl/6J albino for germline transmission and later to a variety of Cre lines on the C57Bl/6 background.

*Foxp3*$^{\mathrm{Thy1.1}}$ (Liston et al., 2008), *Foxp3*$^{\mathrm{cre}}$ (Rubtsov et al., 2008), *Foxp3*$^{\mathrm{BAC}}$ (Petzold et al., 2014), NKp46$^{\mathrm{cre}}$ (Narni-Mancinelli et al., 2011) and CD23$^{\mathrm{cre}}$ (Kwon et al., 2008) mice were used on the C57BL/6 background. *Il2*$^{\mathrm{GFP}}$ mice (Yui et al., 2001) were purchased from MMRRC as stock 009974-MU. *Il2*$^{\mathrm{cre}}$ mice stock 029619 (Yamamoto et al., 2013), CD4$^{\mathrm{cre}}$ stock 022071, CD8$^{\mathrm{cre}}$ stock 008766, *Osx*$^{\mathrm{cre}}$ stock 006361, *Cd19*$^{\mathrm{cre}}$ stock 006785, *H2*$^{dlAb1\text{-}Ea}$ (MHCII$^{-/-}$) stock 003584, *CD1d*$^{-/-}$ stock 008881 and *B2m*$^{-/-}$ stock 002070 mice were purchased from Jackson. *Clec9a*$^{\mathrm{cre}}$ (Schraml et al., 2013) mice were kindly provided by Caetano Reis e Sousa. *Il2*$^{\mathrm{fl/fl}}$ mice (Popmihajlov et al., 2012) were kindly provided by Michael Farrer. *Kb*$^{-/-}$*Db*$^{-/-}$ (MHCI$^{-/-}$) mice were kindly provided by Carla Shatz. All mice were housed under SPF conditions and fed a standard chow diet, *ad libitum*. Mice were assessed at 3-18 weeks of age with littermate controls, unless stated otherwise. C57BL/6.SJL-

*Ptprc*$^{a}$/BoyJ (CD45.1) mice were irradiated with 10 Gy over two doses and reconstituted intravenously with 2-5x10$^{6}$ total BM cells. Mice were left for at least 7 weeks to allow reconstitution before initiating experiments. For neutralization experiments, three-week old littermates were treated with 500 µg anti-IL-5 (BE0198, BioXCell) or isotype control (BE0088, BioXCell) intraperitoneally twice per week for six doses. All experiments were performed in accordance with the University of Leuven Animal Ethics Committee guidelines, the Babraham Institute Animal Welfare and Ethics Review Body or the Animal Care Committee at Maisonneuve-Rosemont Hospital Research Centre. Animal husbandry and experimentation complied with existing European Union and national legislation and local standards, or the Canadian Council on Animal Care guidelines. Sample sizes for mouse experiments were chosen in conjunction with the ethics committees to allow for robust sensitivity without excessive use.

*Flow cytometry*

To obtain single cell suspensions, spleens and lymph nodes were disrupted with glass slides and bones were crushed with a mortar and pestle, before being filtered through 100µm mesh. Lung tissue was digested with 0.4mg/ml Collagenase D (Roche) and 40mg/ml DNase I (Sigma-Aldrich) prepared in RPMI (Invitrogen) supplemented with 2mM $MgCl_2$, 2mM $CaCl_2$, 20% FBS and 2mM HEPES at 37 °C for 30 minutes, followed by filtration through 100 µm mesh. Spleen, bone marrow and lung subsequently underwent red blood cell lysis. Cells were counted using a Countess cell counter (Thermo Fisher). Approximately two million cells were stained with flow cytometry antibodies. Non-specific binding was blocked using 2.4G2 supernatant for mouse cells and dead cells were labelled by fixable viability dye eFluor 780 (ThermoFisher). Cells were fixed and permeabilized with 2% paraformaldehyde or Foxp3 Transcription Factor Staining Buffer Set (eBioscience) according to the manufacturer's instructions. Antibodies used for staining are listed in **Supplementary Table 1**. For cytokine analysis, cells were first stimulated with 500µg/ml phorbol 12,13-dibutyrate (PdBU), 750µg/ml ionomycin and 2µg/ml Brefeldin A (all Tocris Bioscience) in RPMI (Invitrogen) for 4 hours at 37 °C. For phosphoSTAT staining, cells were fixed with 2% paraformaldehyde for 30 min, followed by permeabilisation with ice-cold 100% methanol for 30 min at 4°C. Cells were stained with anti-phosphoSTAT antibodies in PBS with 2.5% FCS and 2mM EDTA overnight at room temperature. Flow cytometry samples were acquired on a

Yeti/ZE5 (Propel Labs/BioRad), Symphony (BD Biosciences), Fortessa (BD Biosciences) or Aurora (Cytek) spectral flow cytometer. Data was compensated via AutoSpill (Roca et al., 2020).

*In vitro cytokine stimulation*

Splenocytes were processed as described above and resuspended in complete RPMI (Invitrogen). Cells were stimulated with 100ng/ml IL-2, IL-7 or IL-15 for 25 min at 37 °C. Cells were immediately fixed with 2% formaldehyde (5% formalin) for phosphoSTAT staining.

*In vitro suppression assay*

$CD4^+$ Treg ($CD4^+CD25^+$) and $CD8^+$ Treg ($CD8^+CD25^+CD103^+$) from *Cd19*$^{cre}$*Rosa*$^{IL\text{-}2}$ and littermate mice and Tconv ($Thy1.1^-CD44^{lo}CD62L^{hi}$) from *Foxp3*$^{Thy1.1}$ mice were isolated from spleens and lymph nodes by negative selection with MagniSort Streptavidin Negative Selection Beads (Thermo Fisher) followed by cell sorting (BD FACSAria III). Antigen-presenting cells were sourced by digesting *Rag*$^{-/-}$ spleens. For the suppression assay, $1x10^5$ Treg were plated per well in complete RPMI along with $5x10^4$ *Rag*$^{-/-}$ splenocytes preincubated with 1 µg/ml anti-CD3 (Thermo Fisher). $CD4^+$ and $CD8^+$ Tconv (responders) were labelled with CellTrace Violet (Thermo Fisher), plated at the indicated ratios with Treg and incubated for 3 days at 37 °C. Suppression was calculated by comparing the proliferative index of the sample with the proliferative index of Tconv cultured in identical conditions except without co-cultured Treg.

*Serum cytokine analysis*

Serum samples were analysed for IL-2, IL-5 and IL-13 levels using ProCartaPlex Immunoassays (Thermo Fisher) and acquired on the Luminex Bio-Plex 3D Suspension Array System (Bio-Rad).

*Immunofluorescence*

Lymph nodes from C57BL/6 mice were embedded in optimal cutting temperature compound (OCT CryoMatrix; Fisher Scientific) and snap frozen in the vapor phase of liquid nitrogen. Ten micrometer sections were generated on a Leica CM1850 Cryostat, then fixed in 2% paraformaldehyde, followed by incubation with 2% Triton X-100 for 30 min and blocking with 5% BSA for 1h at room temperature. Sections were stained overnight with anti-CD8α-eFluor 450 (1:20, 48-0081-82, eBioscience), anti-IgD-Alexa Fluor 594 (1:400, 405740, BioLegend), and anti-Foxp3-Alexa Fluor 488 (1:200, 53-5773-82, Thermofisher) and mounted with Fluormount-G. Confocal images were obtained by using a single-plane confocal microscopy on a Zeiss LSM 780 system with 20x objective. Confocal images were processed in ImageJ (https://imagej.nih.gov/ij/download.html). The total $CD8^+$ cells within the B follicle ($IgD^+$ area) were quantified using the multipoint selection tool when a red or dark center (nucleus) surrounded by CD8 surface staining could be identified. Then, $Foxp3^+$ among $CD8^+$ cells within the B follicle area were quantified.

*Statistics*

All statistical analysis was performed using GraphPad Prism or R. Comparisons between groups were performed using paired or unpaired two-tailed Student's t tests, one-way ANOVA or two-way ANOVA as appropriate. When required, post hoc Sidak's or Tukey's multiple comparison tests were performed. Survival data was analysed using Mantel-Cox log-rank test. Non-parametric testing was performed when data was not normally distributed. FlowSOM, heatmap analysis and PCA were performed in R (version 3.6.2) using in-house scripts. Comparison of tSNE plots and dendograms were performed in R using in-house scripts (Pasciuto et al., 2020). Values are represented as mean ± SEM, unless otherwise indicated.

**Supplementary Figure 1: Technical optimization of Foxp3 and IL-2 co-staining by intracellular flow cytometry.** C57BL/6 splenocytes were stimulated with PdBu (500ng/ml), ionomycin (750ng/ml) and brefeldin A (1μg/ml) for 4hrs to induce IL-2 production and intracellular accumulation. After surface staining for viability and CD antigens, the cells were fixed with either the eBioscience Foxp3/Transcription Factor Staining Buffer Set or using formalin. After fixation, the cells were stained for either 30min or overnight (16hrs) for IL-2 and Foxp3. Representative results demonstrate the increase in IL-2 sensitivity observed with formalin fixation and overnight staining.

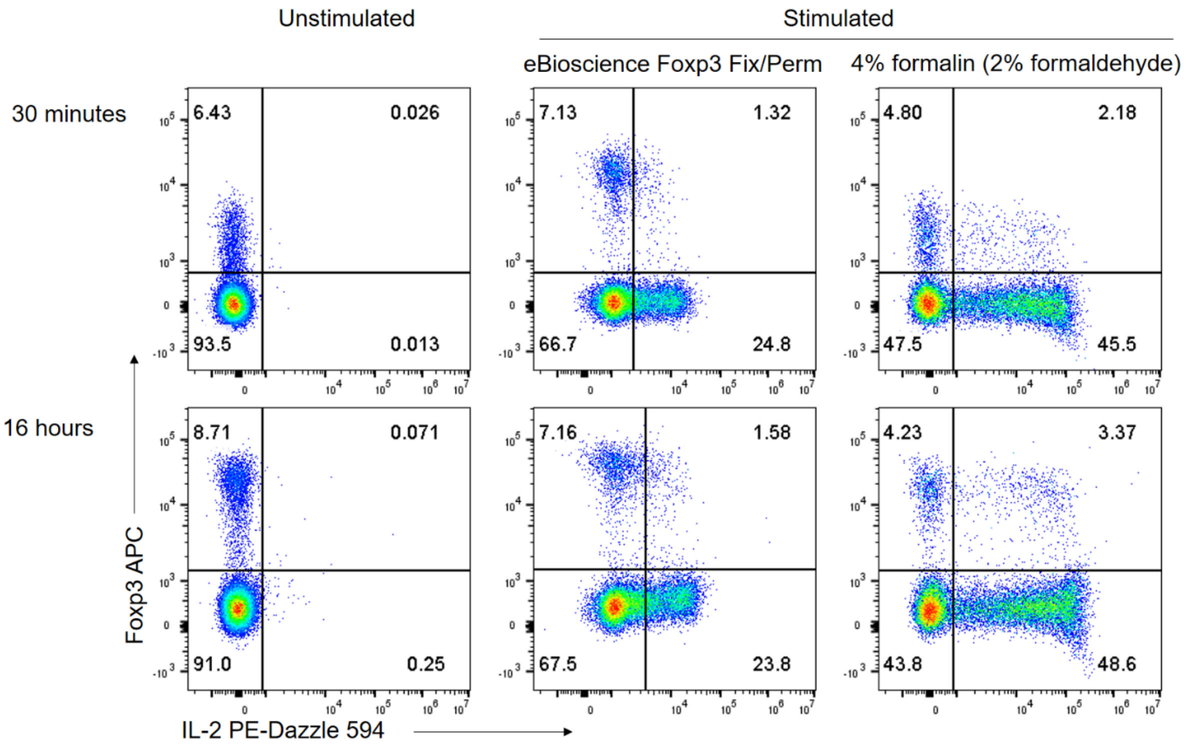

**Supplementary Figure 2. IL-2 reporter expression across leukocyte populations. (A)** IL-2 expression in spleen, LN and lung. n=6. **(B)** Frequency of each annotated cell type among the total IL-2$^+$ population following *ex vivo* stimulation.

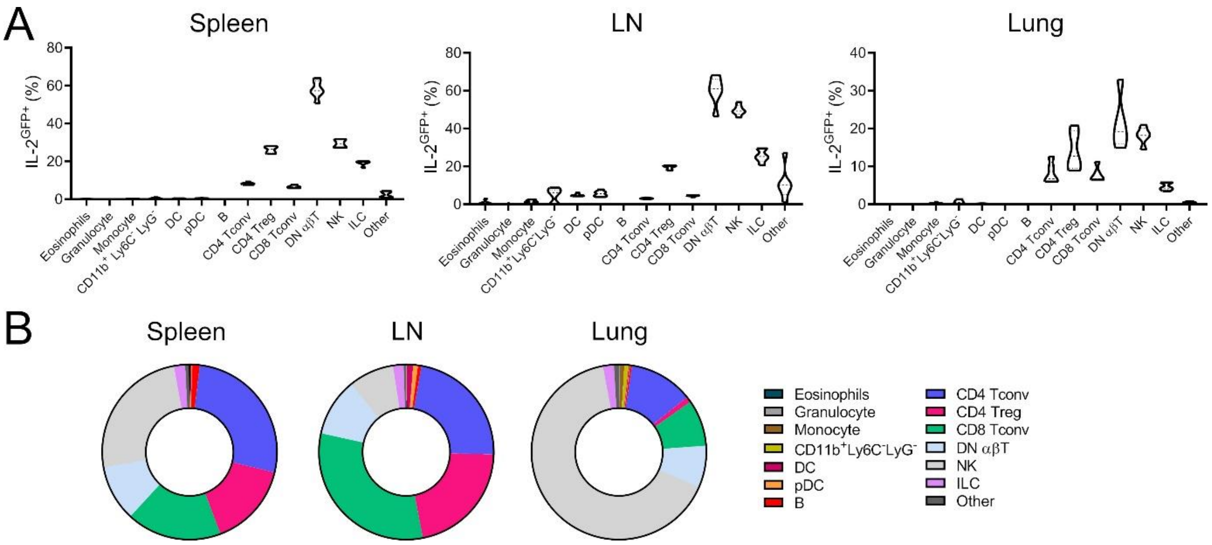

**Supplementary Figure 3. Schematic of Rosa$^{IL\text{-}2}$ mouse.** Construction of targeting vector and Rosa26 integration locus for the generation of Rosa$^{IL\text{-}2}$ mice.

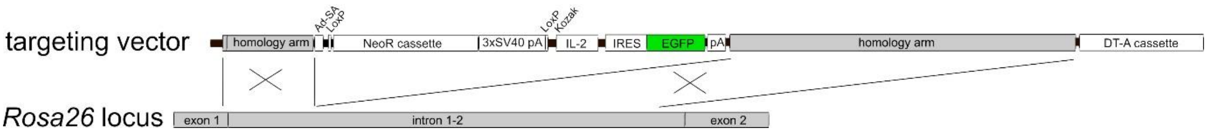

**Supplementary Figure 4. Thymic involution following IL-2 production at the double positive stage. (A)** Thymic cellularity of 4-6-week-old CD8$^{cre}$ *Rosa$^{IL-2}$*, CD4$^{cre}$ *Rosa$^{IL-2}$* or littermate controls. n=12-34. **(B)** Representative gating and **(C)** frequency of thymocyte subsets. n=9-21. **(D)** Frequency of Foxp3$^{+}$ cells among total CD4 T cells. n=9-21. Data pooled from ≥ 2 independent experiments. Significance was tested by one-way ANOVA.

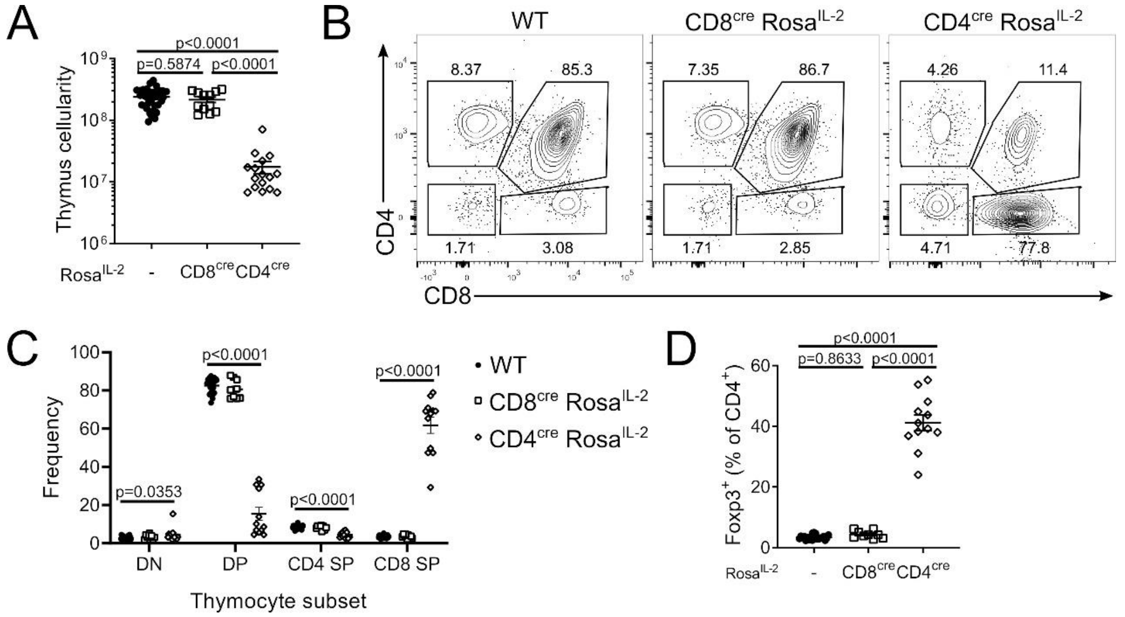

**Supplementary Figure 5. T cell-driven IL-2 induces large-scale phenotypic shifts in CD4 and CD8 T cells. (A)** CD4 T cell data from 4-6-week-old CD8$^{cre}$ Rosa$^{IL2}$, CD4$^{cre}$ Rosa$^{IL2}$ or littermate controls from Figure 2E. Average frequency of each cluster per mouse. **(B)** Dendrogram showing comparative similarity calculated using cross-entropy distributions from tSNE. **(C)** Heat map showing differential marker expression in annotated FlowSOM clusters. **(D)** Heat map showing differential marker expression in total CD4$^{+}$ T cells from different strains, showing mouse replicates. **(E)** CD8 T cell data from 4-6-week-old CD8$^{cre}$ Rosa$^{IL2}$, CD4$^{cre}$ Rosa$^{IL2}$ or littermate controls from Figure 2E. Average frequency of each cluster per mouse. **(F)** Dendrogram showing comparative similarity calculated using cross-entropy distributions from tSNE. **(G)** Heat map of differential marker expression in annotated FlowSOM clusters. **(H)** Heat map of differential marker expression in total CD8$^{+}$ T cells from different strains, showing mouse replicates.

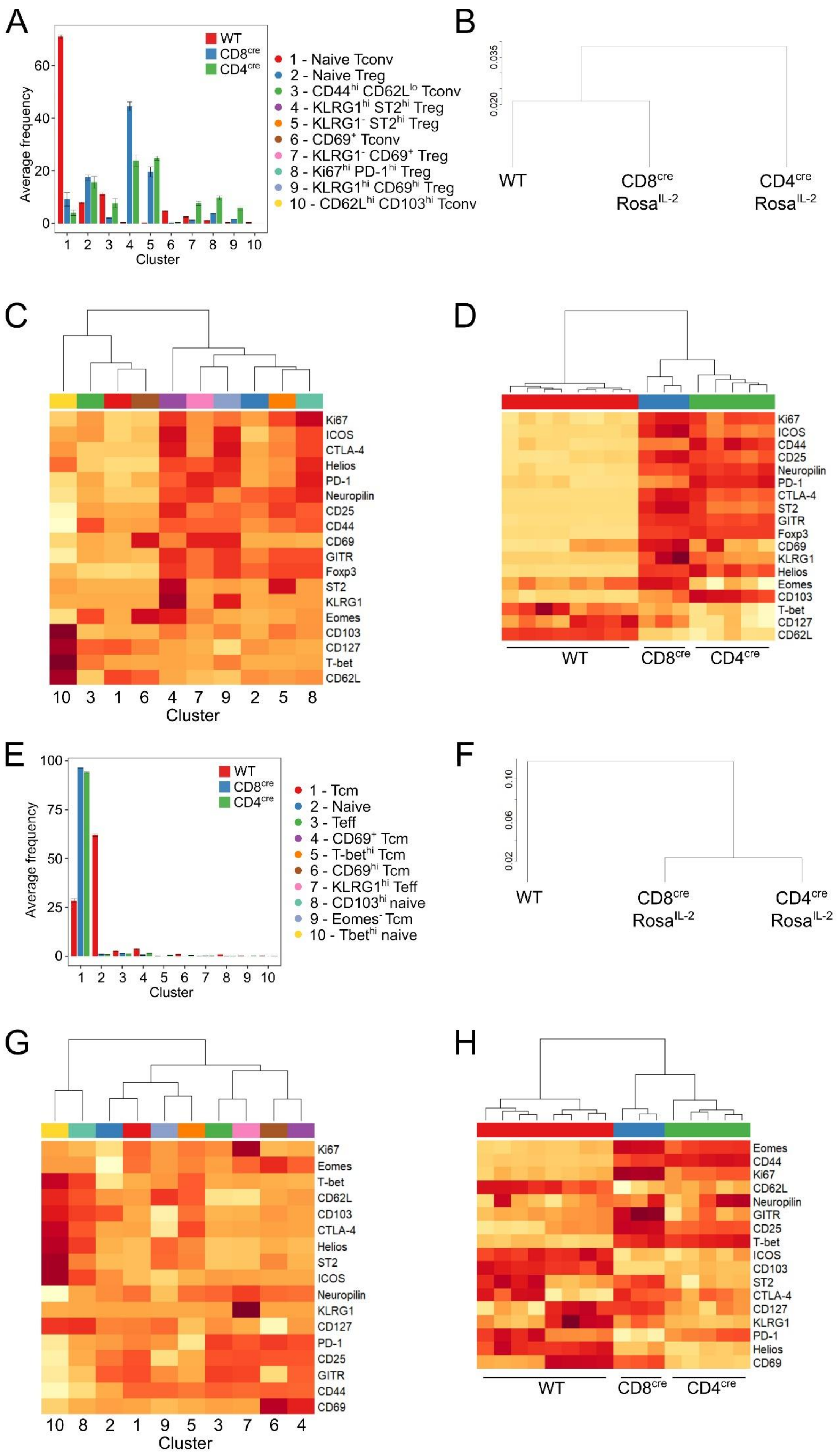
A
WT
CD8$^{cre}$
CD4$^{cre}$
Average frequency
Cluster
1 - Naive Tconv
2 - Naive Treg
3 - CD44$^{hi}$ CD62L$^{lo}$ Tconv
4 - KLRG1$^{hi}$ ST2$^{hi}$ Treg
5 - KLRG1$^{-}$ ST2$^{hi}$ Treg
6 - CD69$^{+}$ Tconv
7 - KLRG1$^{-}$ CD69$^{+}$ Treg
8 - Ki67$^{hi}$ PD-1$^{hi}$ Treg
9 - KLRG1$^{hi}$ CD69$^{hi}$ Treg
10 - CD62L$^{hi}$ CD103$^{hi}$ Tconv
B
WT
CD8$^{cre}$ Rosa$^{IL-2}$
CD4$^{cre}$ Rosa$^{IL-2}$
C
Ki67
ICOS
CTLA-4
Helios
PD-1
Neuropilin
CD25
CD44
CD69
GITR
Foxp3
ST2
KLRG1
Eomes
CD103
CD127
T-bet
CD62L
10 3 1 6 4 7 9 2 5 8
Cluster
D
Ki67
ICOS
CD44
CD25
Neuropilin
PD-1
CTLA-4
ST2
GITR
Foxp3
CD69
KLRG1
Helios
Eomes
CD103
T-bet
CD127
CD62L
WT
CD8$^{cre}$
CD4$^{cre}$
E
WT
CD8$^{cre}$
CD4$^{cre}$
Average frequency
Cluster
1 - Tcm
2 - Naive
3 - Teff
4 - CD69$^{+}$ Tcm
5 - T-bet$^{hi}$ Tcm
6 - CD69$^{hi}$ Tcm
7 - KLRG1$^{hi}$ Teff
8 - CD103$^{hi}$ naive
9 - Eomes$^{-}$ Tcm
10 - Tbet$^{hi}$ naive
F
WT
CD8$^{cre}$ Rosa$^{IL-2}$
CD4$^{cre}$ Rosa$^{IL-2}$
G
Ki67
Eomes
T-bet
CD62L
CD103
CTLA-4
Helios
ST2
ICOS
Neuropilin
KLRG1
CD127
PD-1
CD25
GITR
CD44
CD69
10 8 2 1 9 5 3 7 6 4
Cluster
H
Eomes
CD44
Ki67
CD62L
Neuropilin
GITR
CD25
T-bet
ICOS
CD103
ST2
CTLA-4
CD127
KLRG1
PD-1
Helios
CD69
WT
CD8$^{cre}$
CD4$^{cre}$

**Supplementary Figure 6. IL-2 self-sufficiency drives Treg expansion without altering subset composition.** Splenocyte data from 4-6-week-old *Foxp3*$^{cre/wt}$*Rosa*$^{IL\text{-}2}$, *Foxp3*$^{cre}$*Rosa*$^{IL\text{-}2}$ or littermate controls from Figure 3G. n=5-15. **(A)** Average frequency of each cluster per mouse. **(B)** Heat map showing differential marker expression in annotated FlowSOM clusters. **(C)** Average frequency of each Treg sub-cluster per mouse, from Figure 3J. **(D)** Heat map showing differential marker expression in annotated FlowSOM Treg sub-clusters.

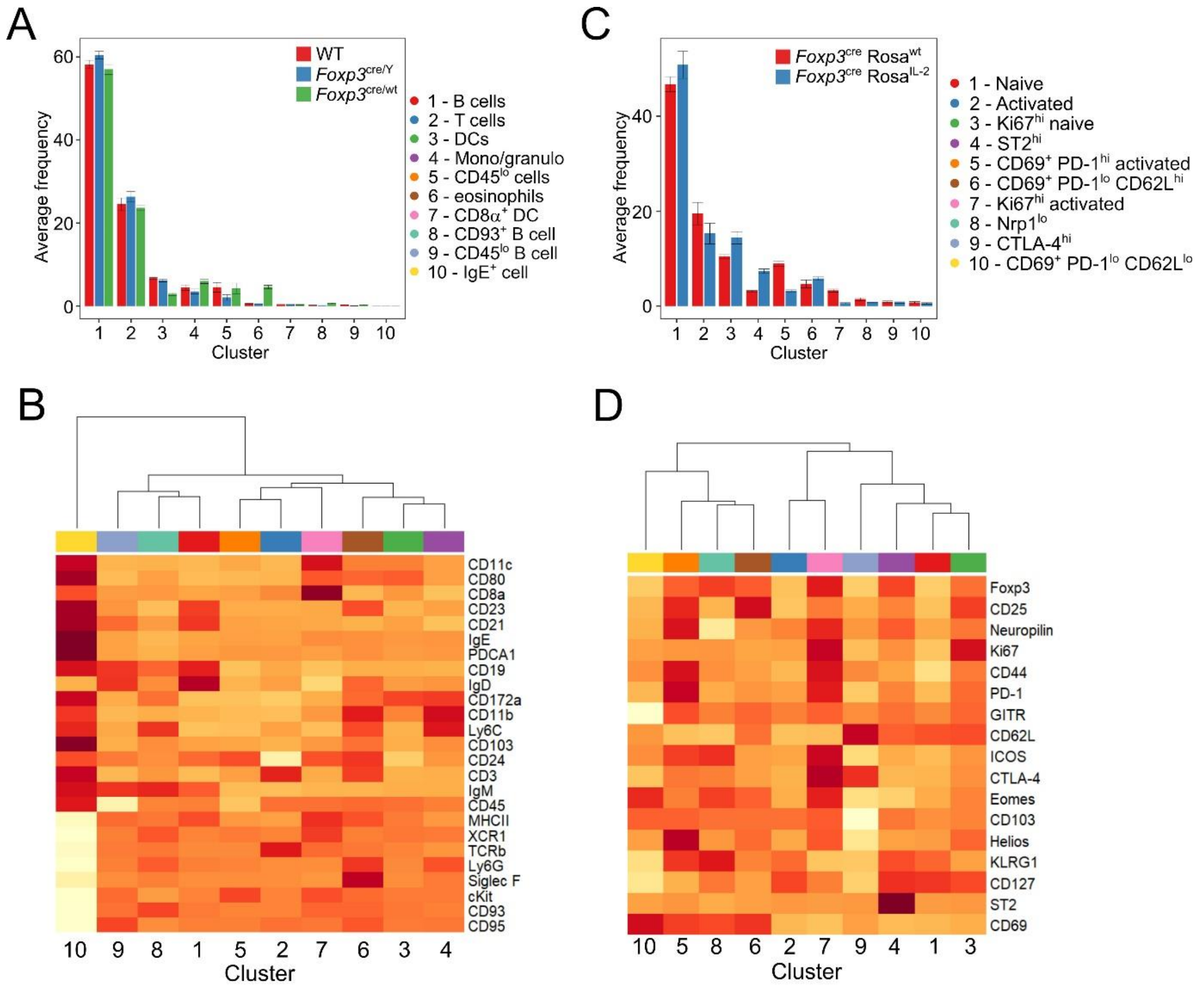

**Supplementary Figure 7. Autocrine IL-2 production drives divergent responses in CD4$^+$ and CD8$^+$ T cells.** Chimeric mice were generated using bone marrow from WT (CD45.1$^+$) and CD4$^{cre}$Rosa$^{IL-2}$ (CD45.2$^+$) at varying ratios. n=4-6, pooled from two independent experiments. **(A)** Frequency of thymocyte subsets derived from CD4$^{cre}$Rosa$^{IL-2}$ (CD45.2$^+$) after reconstitution. **(B)** Total T cell numbers in spleen. **(C)** Frequency of naïve (CD44$^{lo}$ CD62L$^{hi}$), $T_{eff}$ (CD44$^{hi}$ CD62L$^{lo}$) or $T_{cm}$ (CD44$^{hi}$ CD62L$^{hi}$) derived from CD4$^{cre}$Rosa$^{IL-2}$ (CD45.2$^+$) bone-marrow in the spleen. **(D)** Comparison of splenocyte frequencies from experimental chimeras (50% WT, 50% CD4$^{cre}$Rosa$^{IL-2}$) with control chimeras (50% WT, 50% CD4$^{cre}$Rosa$^{wt}$) generated simultaneously. **(E)** Heat map showing differential marker expression in annotated FlowSOM clusters from Figure 4E for CD4$^+$ T cells or **(F)** CD8$^+$ T cells.

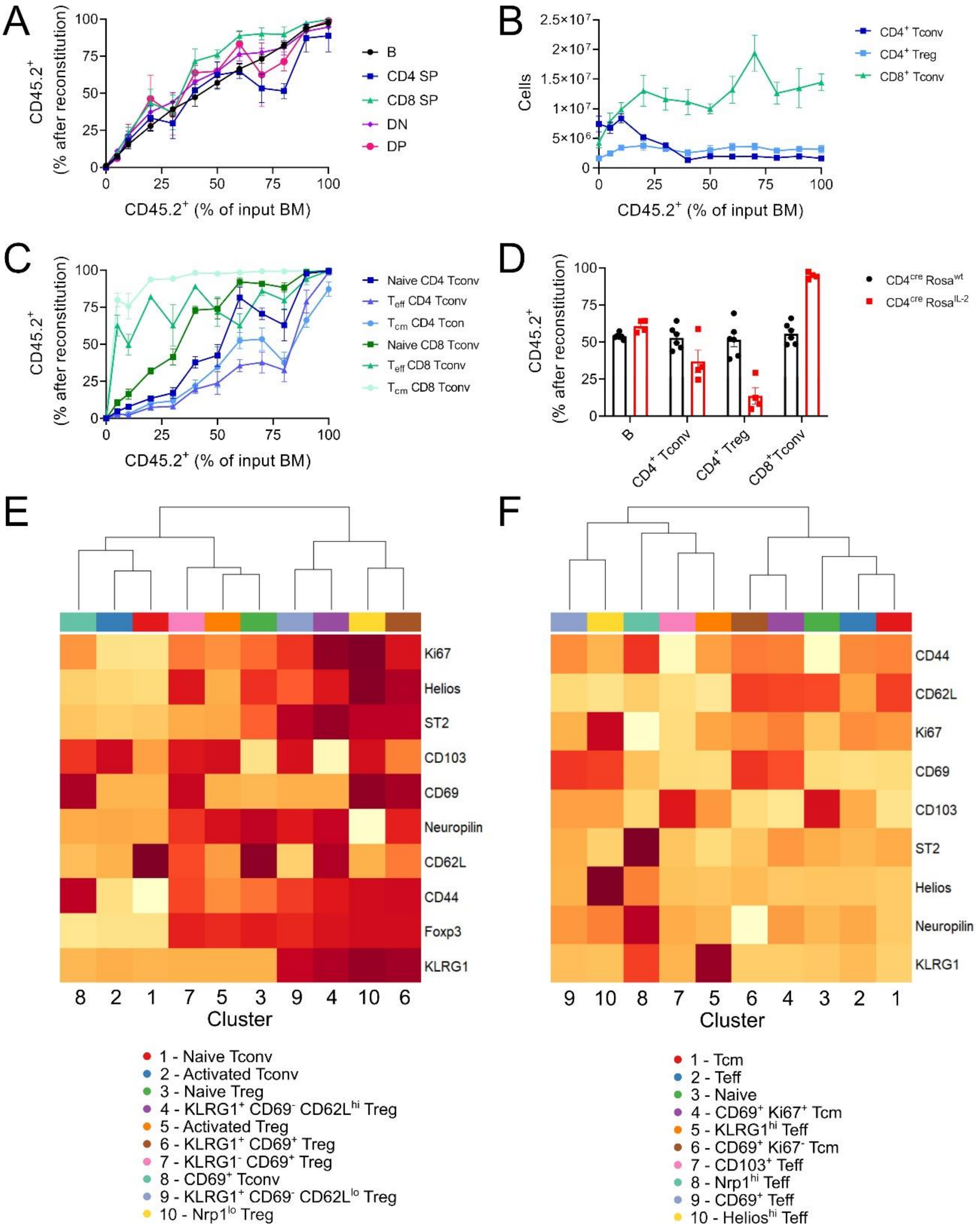
A
CD45.2+ (% after reconstitution)
CD45.2+ (% of input BM)
B
CD4 SP
CD8 SP
DN
DP
B
Cells
CD45.2+ (% of input BM)
CD4+ Tconv
CD4+ Treg
CD8+ Tconv
C
CD45.2+ (% after reconstitution)
CD45.2+ (% of input BM)
Naive CD4 Tconv
Teff CD4 Tconv
Tcm CD4 Tcon
Naive CD8 Tconv
Teff CD8 Tconv
Tcm CD8 Tconv
D
CD45.2+ (% after reconstitution)
CD4cre Rosawt
CD4cre RosaIL-2
B
CD4+ Tconv
CD4+ Treg
CD8+Tconv
E
Ki67
Helios
ST2
CD103
CD69
Neuropilin
CD62L
CD44
Foxp3
KLRG1
8 2 1 7 5 3 9 4 10 6
Cluster
1 - Naive Tconv
2 - Activated Tconv
3 - Naive Treg
4 - KLRG1+ CD69- CD62Lhi Treg
5 - Activated Treg
6 - KLRG1+ CD69+ Treg
7 - KLRG1- CD69+ Treg
8 - CD69+ Tconv
9 - KLRG1+ CD69- CD62Llo Treg
10 - Nrp1lo Treg
F
CD44
CD62L
Ki67
CD69
CD103
ST2
Helios
Neuropilin
KLRG1
9 10 8 7 5 6 4 3 2 1
Cluster
1 - Tcm
2 - Teff
3 - Naive
4 - CD69+ Ki67+ Tcm
5 - KLRG1hi Teff
6 - CD69+ Ki67- Tcm
7 - CD103+ Teff
8 - Nrp1hi Teff
9 - CD69+ Teff
10 - Helioshi Teff

**Supplementary Figure 8. IL-2 production alters cytokine responses in CD4$^+$ and CD8$^+$ T cells. (A)** MFI and **(B)** percent of cells expressing CD25 (IL-2Rα) on CD4$^+$ Treg, CD4$^+$ Tconv and CD8$^+$ T cells in WT and CD4$^{Cre}$ Rosa$^{IL-2}$ transgenic mice. (**C)** Ratio of CD25 MFI to CD132 (common γ chain) MFI. **(D)** Upregulation of pSTAT3 in response to cytokine stimulation in CD4$^+$ Treg, naïve CD4$^+$ Tconv and CD8$^+$ T cells. Significance was tested by Sidak's multiple comparison test on 2-way ANOVA.

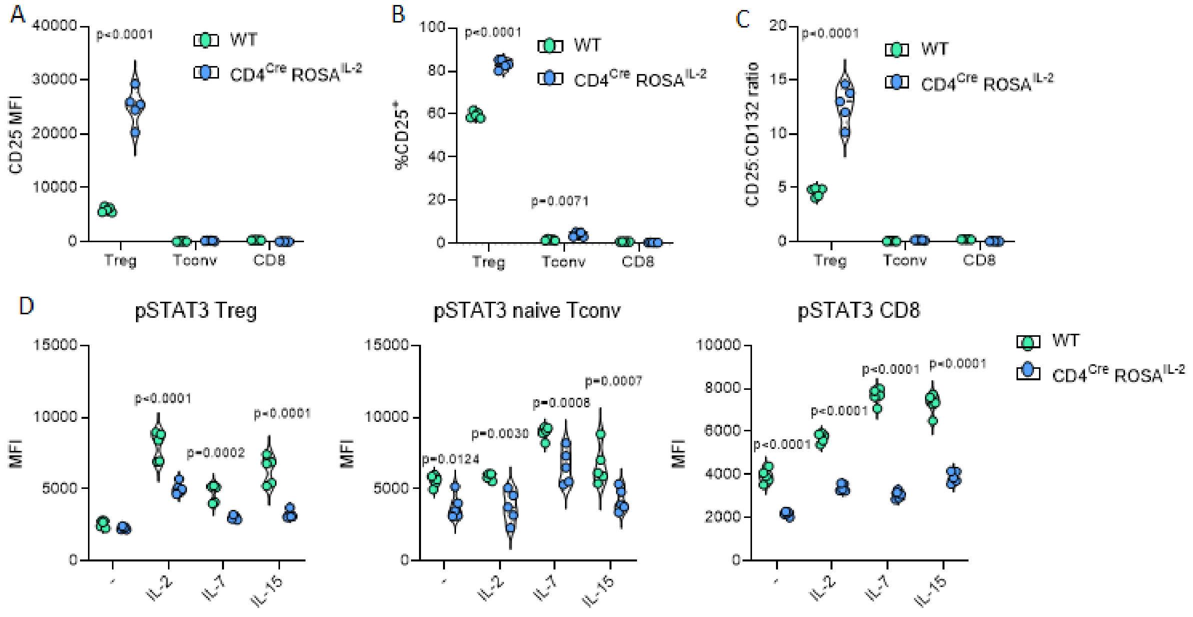

**Supplementary Figure 9. Dendritic cell-driven IL-2 favors Treg expansion. (A)** Cellularity of spleen and LN of 4-6-week-old *Clec9a*$^{cre}$ *Rosa*$^{IL\text{-}2}$ and littermate controls. **(B)** Frequency of GFP expression among DC subsets. **(C)** Representative gating, frequency and number of splenic DCs. **(D)** Frequency of CD8a$^{+}$, CD11b$^{+}$ and CD103$^{+}$ splenic DCs. **(E)** Frequency and number of CD4$^{+}$ Tconv, **(F)** CD8$^{+}$ Tconv and **(G)** CD4$^{+}$ Treg in spleen. **(H)** Frequency of Foxp3$^{+}$ cells among total CD4 T cells. **(I)** Representative tSNE of high-parameter flow cytometry data from splenic CD4$^{+}$ (top) and CD8$^{+}$ (bottom). **(J)** Average frequency of each cluster per mouse for CD4 T cells or **(K)** CD8 T cells. **(L)** Frequency of effector (CD44$^{hi}$ CD62L$^{lo}$) or central memory (CD44$^{hi}$ CD62L$^{hi}$) cells from CD4 Tconv or **(M)** CD8 Tconv. Data pooled from 3 independent experiments with 8-10 mice per genotype. Significance was tested by unpaired t-test (A-H, L, M) or two-sample Kolmogorov-Smirnov test (I).

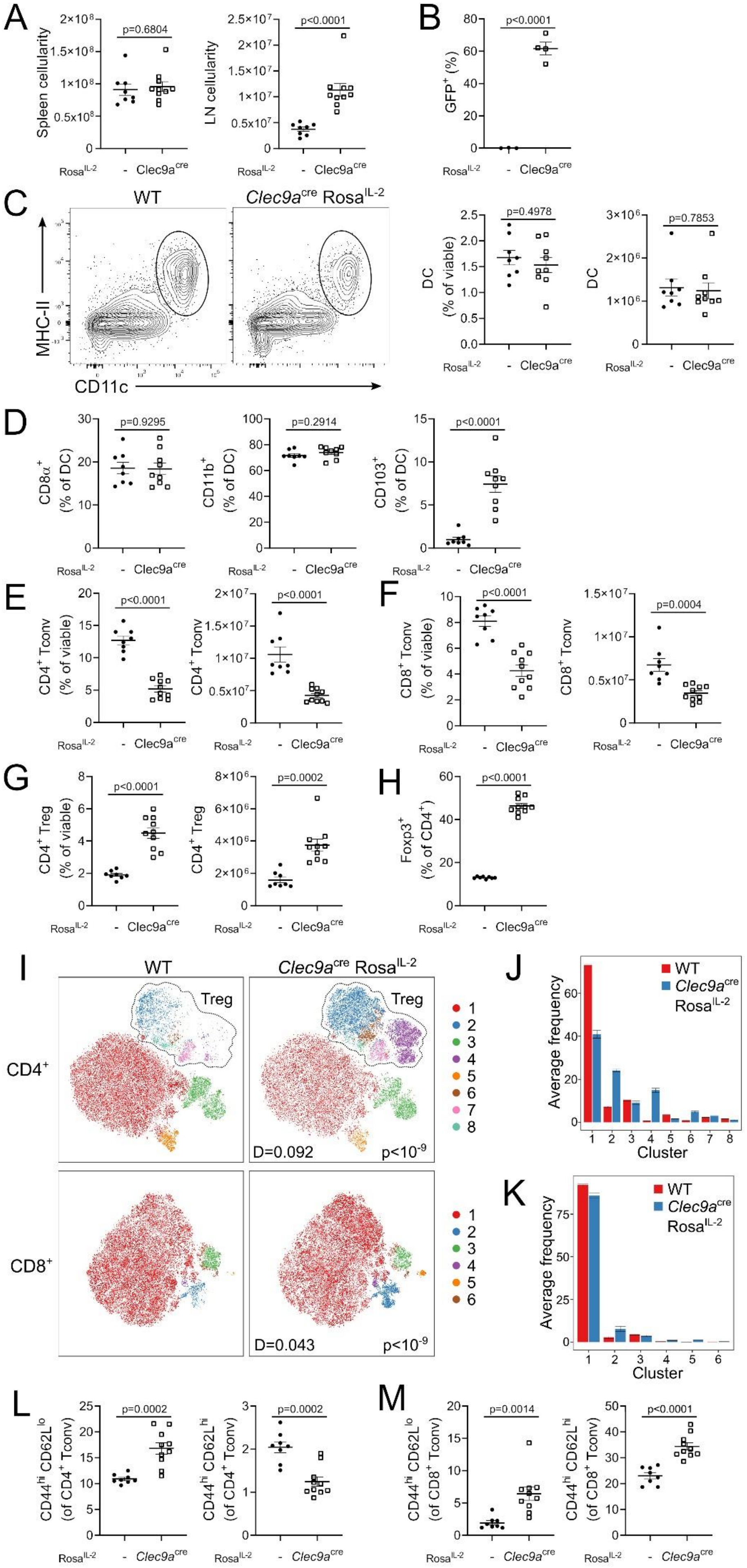

A
Spleen cellularity
p=0.6804
LN cellularity
p<0.0001
B
GFP+ (%)
p<0.0001
Rosa IL-2
-
Clec9acre
C
WT
Clec9acre RosaIL-2
MHC-II
CD11c
DC (% of viable)
p=0.4978
DC
p=0.7853
D
CD8α+ (% of DC)
p=0.9295
CD11b+ (% of DC)
p=0.2914
CD103+ (% of DC)
p<0.0001
E
CD4+ Tconv (% of viable)
p<0.0001
CD4+ Tconv
p<0.0001
F
CD8+ Tconv (% of viable)
p<0.0001
CD8+ Tconv
p=0.0004
G
CD4+ Treg (% of viable)
p<0.0001
CD4+ Treg
p=0.0002
H
Foxp3+ (% of CD4+)
p<0.0001
I
WT
Clec9acre RosaIL-2
Treg
CD4+
CD8+
D=0.092
p<10-9
D=0.043
p<10-9
J
WT
Clec9acre RosaIL-2
Average frequency
Cluster
K
WT
Clec9acre RosaIL-2
Average frequency
Cluster
L
CD44hi CD62Llo (of CD4+ Tconv)
p=0.0002
CD44hi CD62Lhi (of CD4+ Tconv)
p=0.0002
M
CD44hi CD62Llo (of CD8+ Tconv)
p=0.0014
CD44hi CD62Lhi (of CD8+ Tconv)
p<0.0001

**Supplementary Figure 10. NK cell-driven IL-2 selectively favors NK expansion. (A)** Cellularity of spleen and LN of 4-6-week-old NKp46$^{cre}$ *Rosa*$^{IL\text{-}2}$ and littermate controls. **(B)** Frequency of GFP expression among NK cells. **(C)** Representative gating, frequency and number of splenic NK cells. **(D)** Frequency and number of CD4 Tconv, **(E)** CD8 Tconv and **(F)** CD4 Treg in spleen. **(G)** Frequency of Foxp3$^{+}$ cells among total CD4$^{+}$ T cells. **(H)** Representative tSNE of high-parameter flow cytometry data from splenic CD4$^{+}$ (top) and CD8$^{+}$ (bottom). **(I)** Average frequency of each cluster per mouse for CD4 T cells or **(J)** CD8 T cells. **(K)** Frequency of effector (CD44$^{hi}$ CD62L$^{lo}$) or central memory (CD44$^{hi}$ CD62L$^{hi}$) cells from CD4 Tconv or **(L)** CD8 Tconv. Data pooled from 2 independent experiments with 4-6 mice per genotype. Significance was tested by unpaired t-test (A-G, K, L) or two-sample Kolmogorov-Smirnov test (H).

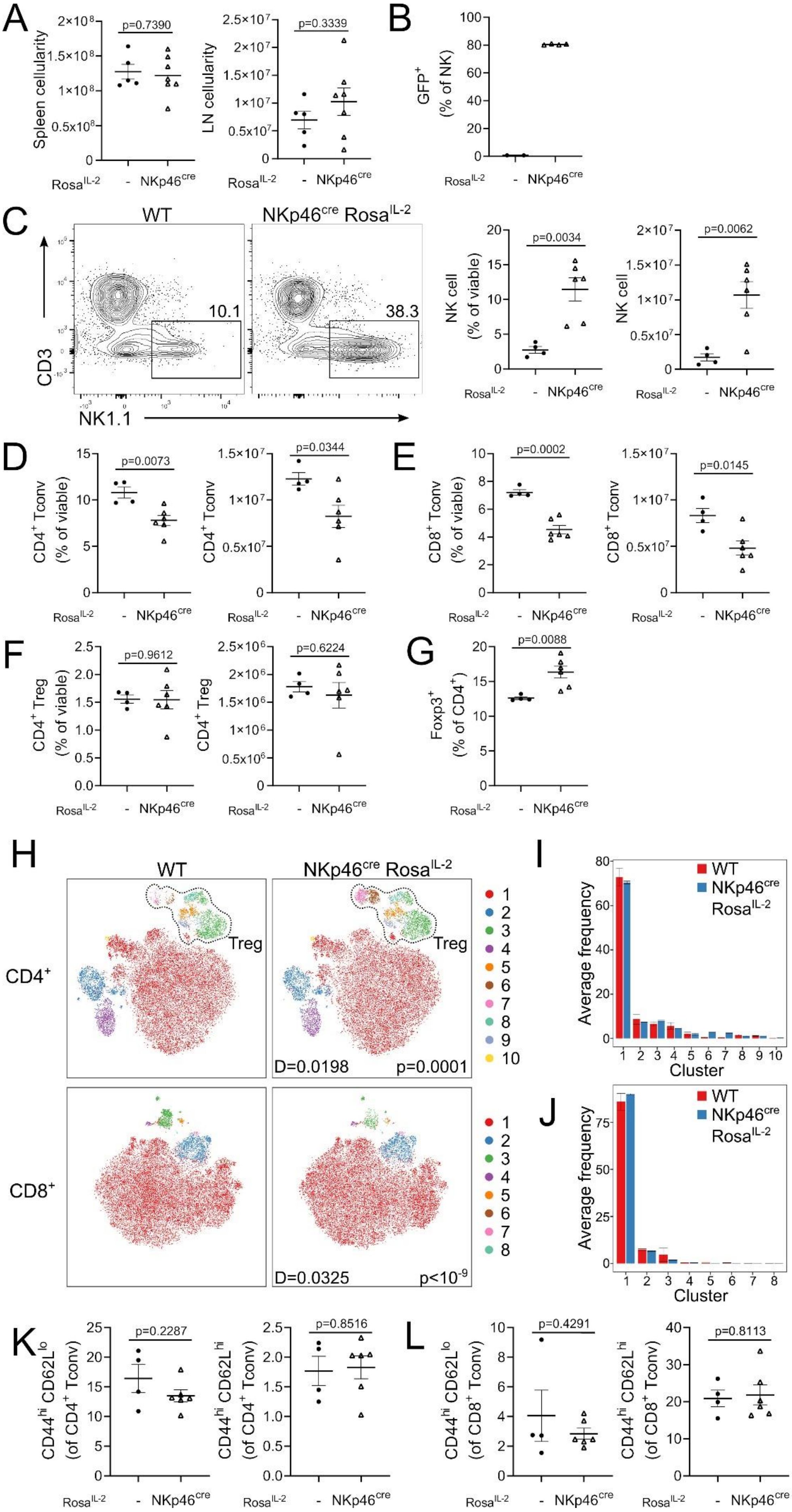

A
Spleen cellularity
p=0.7390
LN cellularity
p=0.3339
Rosa^IL-2 - NKp46^cre
B
GFP+ (% of NK)
C
WT
NKp46^cre Rosa^IL-2
10.1
38.3
CD3
NK1.1
NK cell (% of viable)
p=0.0034
NK cell
p=0.0062
D
CD4+ Tconv (% of viable)
p=0.0073
CD4+ Tconv
p=0.0344
E
CD8+ Tconv (% of viable)
p=0.0002
CD8+ Tconv
p=0.0145
F
CD4+ Treg (% of viable)
p=0.9612
CD4+ Treg
p=0.6224
G
Foxp3+ (% of CD4+)
p=0.0088
H
WT
NKp46^cre Rosa^IL-2
CD4+
Treg
D=0.0198
p=0.0001
CD8+
D=0.0325
p<10^-9
I
Average frequency
WT
NKp46^cre Rosa^IL-2
Cluster
J
Average frequency
WT
NKp46^cre Rosa^IL-2
Cluster
K
CD44hi CD62Llo (of CD4+ Tconv)
p=0.2287
CD44hi CD62Lhi (of CD4+ Tconv)
p=0.8516
L
CD44hi CD62Llo (of CD8+ Tconv)
p=0.4291
CD44hi CD62Lhi (of CD8+ Tconv)
p=0.8113

**Supplementary Figure 11. Expression of IL-2 by B cells drives an ILC2-eosinophil-orientated cellular circuit in the bone-marrow.** **(A)** Bone-marrow cellularity of 4-6-week-old *Cd19*$^{cre}$ *Rosa*$^{IL\text{-}2}$ mice and littermate controls. n=9-11. **(B)** Frequency of eosinophils in the bone-marrow. n=6-14. **(C)** Representative gating and frequency of myeloid progenitors (MP; lineage$^{-}$ Sca-1$^{-}$ c-kit$^{+}$) cells in the bone-marrow, pre-gated on CD45+ lineage$^{-}$. n=3. **(D)** Representative gating and frequency of GMPs (granulocyte-monocyte progenitor) in the bone-marrow (pre-gated from MP gate). n=3. **(E)** IL-5 expression among cellular lineages in the bone-marrow. n=6-9. **(F)** Frequency of GATA3$^{+}$ ILC2 in BM. n=4-6. **(G)** Frequency of eosinophils in the bone-marrow of mice treated with anti-IL-5 neutralising antibody. n=3-8 mice. Significance was tested by unpaired t-test (A, C-F) or one-way ANOVA (B, G).

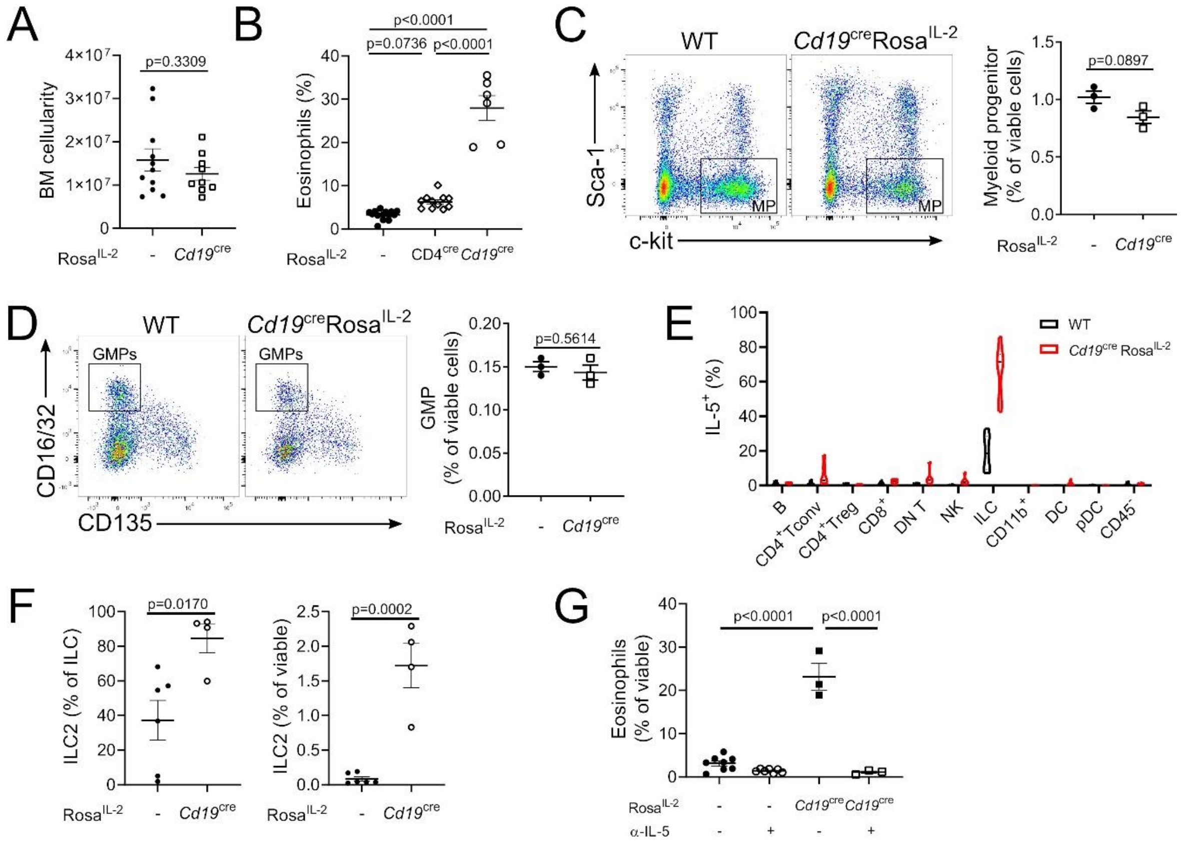
A
p=0.3309
BM cellularity
Rosa^IL-2 - Cd19^cre
B
p<0.0001
p=0.0736
p<0.0001
Eosinophils (%)
Rosa^IL-2 - CD4^cre Cd19^cre
C
WT
Cd19^cre Rosa^IL-2
Sca-1
c-kit
MP
p=0.0897
Myeloid progenitor (% of viable cells)
Rosa^IL-2 - Cd19^cre
D
WT
Cd19^cre Rosa^IL-2
GMPs
CD16/32
CD135
p=0.5614
GMP (% of viable cells)
Rosa^IL-2 - Cd19^cre
E
WT
Cd19^cre Rosa^IL-2
IL-5^+ (%)
B
CD4^+ Tconv
CD4^+ Treg
CD8^+
DN T
NK
ILC
CD11b^+
DC
pDC
CD45^-
F
p=0.0170
ILC2 (% of ILC)
Rosa^IL-2 - Cd19^cre
p=0.0002
ILC2 (% of viable)
Rosa^IL-2 - Cd19^cre
G
p<0.0001
p<0.0001
Eosinophils (% of viable)
Rosa^IL-2 - - Cd19^cre Cd19^cre
α-IL-5 - + - +

**Supplementary Figure 12. Expression of IL-2 by mature B cells drives the ILC2-eosinophil circuit. (A)** Spleen weight and cellularity of 4-6-week-old *CD23*$^{cre}$ *Rosa*$^{IL\text{-}2}$ mice and littermate controls. **(B)** Representative tSNE of high-parameter flow cytometry data from splenic viable cells and average frequency of each cluster per mouse. **(C)** Representative gating and frequency of eosinophils in spleen and bone-marrow. **(D)** Frequency of GATA3$^+$ ILC2 among total ILCs in spleen. **(E)** Frequency of ILC2 in spleen. Data pooled from two independent experiments, n=6. Significance was tested by unpaired t-test (A, C-E) or two-sample Kolmogorov-Smirnov test (B).

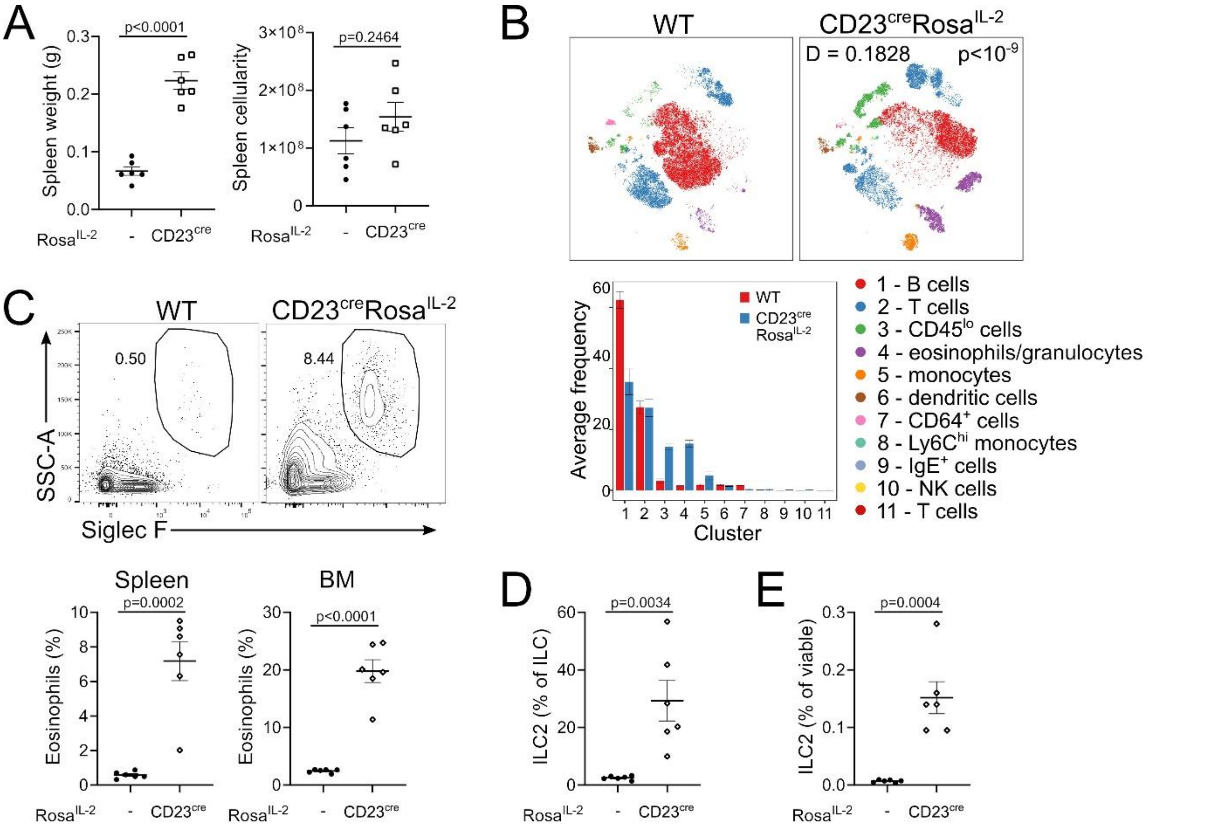

**Supplementary Figure 13. Bone-marrow-localized IL-2 expression does not recapitulate the ILC2-eosinophil circuit. (A)** Spleen weight and cellularity of 4-6-week-old *Osx*$^{cre}$ *Rosa*$^{IL\text{-}2}$ and controls. **(B)** Representative tSNE of high-parameter flow cytometry data from splenic viable cells. **(C)** Average frequency of each cluster per mouse. **(D)** Frequency of eosinophils in spleen and bone-marrow. **(E)** Frequency of GATA3$^{+}$ ILC2 among total ILCs in spleen. **(F)** Frequency of ILC2 in spleen. n=3. Significance was tested by unpaired t-test (A, D-F) or two-sample Kolmogorov-Smirnov test (B).

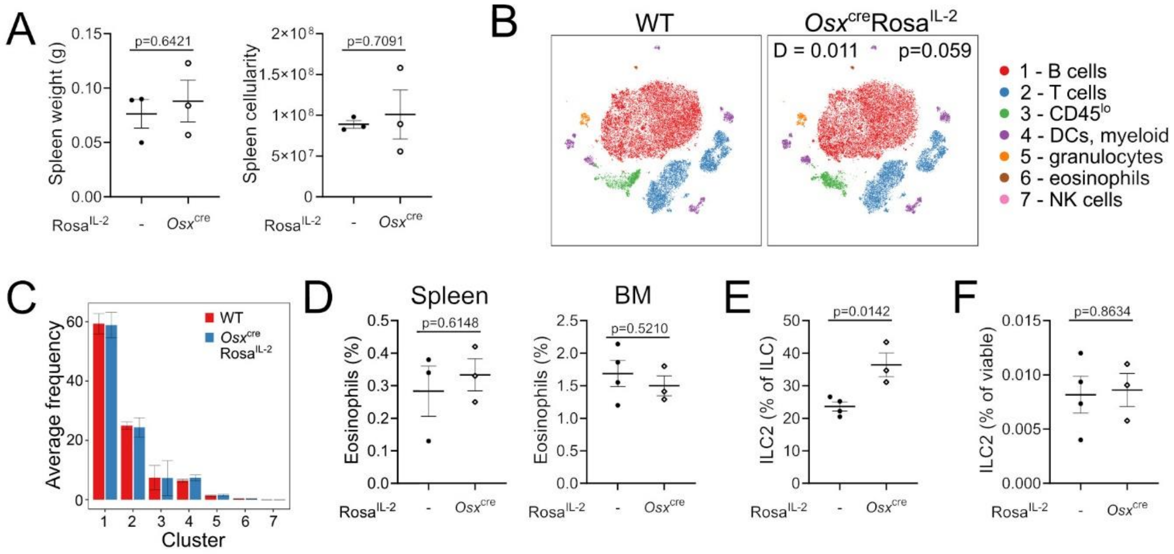

**Supplementary Figure 14. Foxp3 expression in the CD8 lineage. (A)** Hierarchical gating of CD8α$^+$Foxp3$^+$ cells in *Cd19*$^{cre}$Rosa$^{IL-2}$ and littermate controls, pre-gated on viable lymphocytes. **(B)** Expression of the *Foxp3*$^{Thy1.1}$ reporter, **(C)** *Foxp3*$^{YFP-cre}$ reporter and **(D)** Foxp3$^{BAC}$ reporter in CD8α$^+$ CD4$^-$ T cells. **(E)** tSNE representation of high-parameter flow cytometry data from splenic T cell populations from *Cd19*$^{cre}$Rosa$^{IL-2}$ mice and littermate controls. FlowSOM clusters annotated based on differential expression of key markers. **(F)** Expression of CD8β in CD8α$^+$Foxp3$^+$ cells from the spleen of wildtype mice. **(G)** Usage of E8I enhancer (RFP$^+$) in CD8$^+$ T cells from CD8$^{cre}$Rosa$^{RFP}$ mice. n=4, unpaired t-test.s

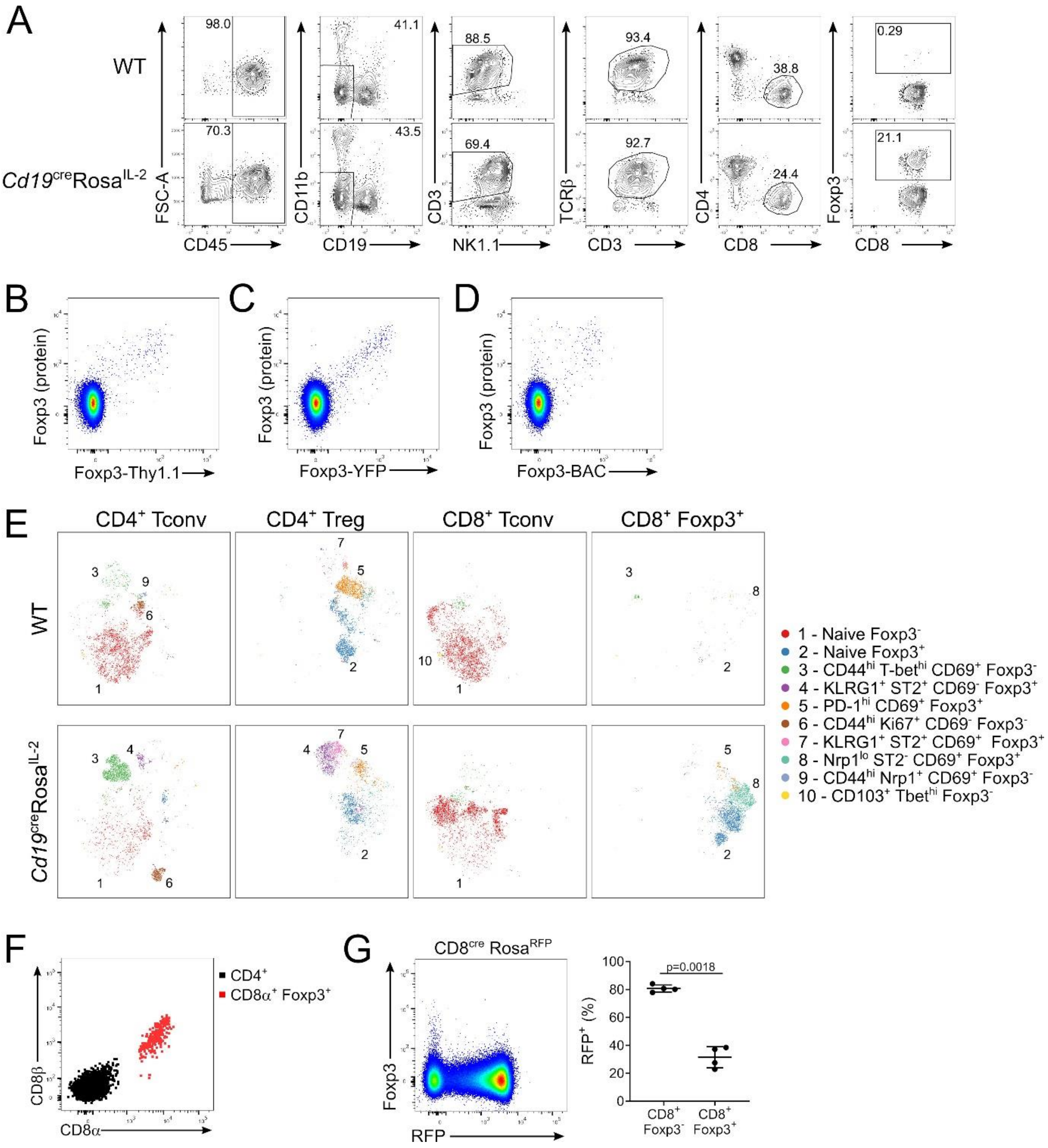

**Supplementary Figure 15. CD8$^+$ Treg are MHC I-restricted. (A)** Representative gating of CD8$^+$Foxp3$^+$ cells in spleens of wildtype, *MHCI$^{-/-}$*, *MHCII$^{-/-}$*, *MHCI$^{-/-}$.MHCII$^{-/-}$* and *CD1d$^{-/-}$* mice. **(B)** Number of CD8$^+$Foxp3$^+$ cells in spleen. **(C)** Frequency and number of CD8$^+$Foxp3$^+$ cells in thymus. n=3-5, data pooled from two independent experiments. Significance tested by one-way ANOVA.

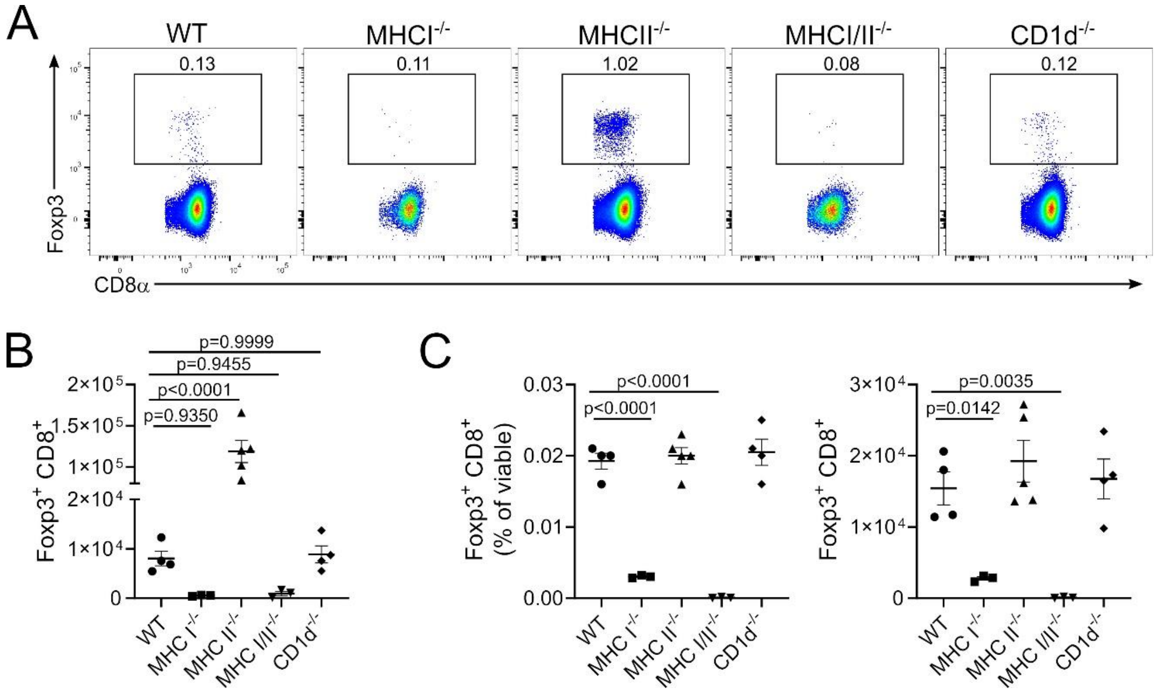
