## Supplementary Table 1 for "Context-dependent effects of IL-2 rewire immunity into distinct cellular circuits"

| Target | Antibody clone | Fluorophore | Identifier | Company |
|---|---|---|---|---|
| B220 | RA3-6B2, rat IgG2a | PE-Cy5 | Catalog number: 15-0452-82 RRID:AB_468755 | eBioscience |
| Bcl-6 | K112-91, mouse IgG1 | AF647 | Catalog number: 561525 RRID:AB_10898007 | BD Biosciences |
| CCR2 | 475301, rat IgG2b | BV480 | Catalog number: 747971 RRID:AB_2872432 | BD Biosciences |
| CCR3 | J073E5, rat IgG2a | AF647 | Catalog number: 144508 RRID:AB_2561536 | BioLegend |
| CCR3 | 83101, rat IgG2a | PerCP | Catalog number: FAB729C-025 | R&D Systems (Biotechne) |
| CCR9 | CW-1.2, mouse IgG2a | BB515 | Catalog number: 565577 RRID:AB_2869688 | BD Biosciences |
| CD103 | 2E7, armenian hamster | BV421 | Catalog number: 121422 RRID:AB_2562901 | BioLegend |
| CD103 | M290, rat IgG2a | BUV395 | Catalog number: 740238 RRID:AB_2739985 | BD Biosciences |
| CD11b | M1/70, rat IgG2b | APC-Cy5.5 | Catalog number: 14-0112-82 RRID:AB_467108 | eBioscience |
| CD11b | M1/70, rat IgG2b | BV750 | Special order | BD Biosciences |
| CD11b | M1/70, rat IgG2b | eFluor450 | Catalog number: 48-0112-82 RRID:AB_1582236 | eBioscience |
| CD11b | M1/70, rat IgG2b | PE-Cy5 | Catalog number: 15-0112-81 RRID:AB_468713 | eBioscience |
| CD11c | N418, armenian hamster | AF405 | Catalog number: FAB69501V-100UG | R&D Systems (Biotechne) |
| CD11c | N418, armenian hamster | BV421 | Catalog number: 117330 RRID:AB_11219593 | BioLegend |
| CD11c | N418, armenian hamster | BV605 | Catalog number: 117334 RRID:AB_2562415 | BioLegend |
| CD117/c-kit | 2B8, rat IgG2b | BUV661 | Special order | BD Biosciences |
| CD122 | TM-b1, rat IgG2b | PE | Catalog number: 12-1222-82 RRID:AB_465836 | eBioscience |
| CD122 | TM-b1, rat IgG2b | PE-Cy5 | Catalog number 123219, RRID:AB_2715961 | BioLegend |
| CD127 | SB/199, rat IgG2b | BUV661 | Catalog number MBS421132, RRID:AB_2083704 | BD Biosciences |
| CD127 | SB/199, rat IgG2b | BUV737 | Catalog number: 564399, RRID:AB_2738791 | BD Biosciences |
| CD127 | A7R34 | PE | Catalog number: 2-1271-82 RRID:AB_2127182 | eBioscience |
| CD132 | Tugm2, rat IgG2b | APC | Catalog number: 132308, RRID:AB_10643576 | BioLegend |
| CD138 | 281-2, rat IgG2a | BV785 | Catalog number: 142534 RRID:AB_2814047 | BioLegend |
| CD19 | 1D3, rat IgG2a | APC | Cat# 17-0193-82, RRID:AB_1659676 | eBioscience |
| CD19 | 1D3, rat IgG2a | BUV661 | Catalog number: 565076 RRID:AB_2739055 | BD Biosciences |
| CD19 | 1D3, rat IgG2a | SuperBright780 | Catalog number: 78-0193-82 RRID:AB_2722936 | eBioscience |
| CD21/35 | 7G6, rat IgG2b | BUV737 | Catalog number: 565090 RRID:AB_2739061 | BD Biosciences |
| CD23 | B3B4, rat IgG2a | PE-Cy7 | Catalog number: 25-0232-82 RRID:AB_469604 | eBioscience |
| CD25 | PC61, rat IgG1 | PE-Cy5.5 | Catalog number: 35-0251-80 RRID:AB_11218285 | eBioscience |
| CD25 | PC61, rat IgG1 | BV421 | Catalog number: 102043 RRID:AB_2562611 | BioLegend |
| CD25 | PC61, rat IgG1 | BV480 | Catalog number: 566120, RRID:AB_2739522 | BD Biosciences |
| CD25 | PC61, rat IgG1 | BV650 | Catalog number: 102038 RRID:AB_2563060 | BioLegend |
| CD3 | 145-2C11, armenian hamster | Biotin | Cat# 13-0032-82, RRID:AB_2572762 | ThermoFisher |
| CD3 | 145-2C11, armenian hamster | AF488 | Catalog number: 100321 RRID:AB_389300 | BioLegend |
| CD3 | 17A2, rat IgG2b | BV650 | Catalog number: 100229 RRID AB_11204249 | BioLegend |
| CD3 | 145-2C11, armenian hamster | PE-Cy7 | Catalog number: 25-0031-82, RRID:AB_469572 | ThermoFisher |
| CD3 | 17A2, rat IgG2b | Spark Blue 550 | Catalog number: 100259, RRID:AB_2819767 | BioLegend |
| CD4 | RM4-5, rat IgG2a | BV605 | Catalog number: 100547, RRID:AB_11125962 | BioLegend |
| CD4 | RM4-5, rat IgG2a | Qdot800 | Catalog number: Q22165 RRID:AB_2556521 | ThermoFisher |
| CD4 | GK1.5, rat IgG2b | BUV496 | Catalog numbere: 564667 RRID:AB_2722549 | BD Biosciences |
| CD4 | GK1.5, rat IgG2b | BUV395 | Catalog number: 563790, RRID:AB_2738426 | BD Biosciences |
| CD44 | IM7, rat IgG2b | BV510 | Catalog number: 103044 RRID:AB_2650923 | BioLegend |
| CD44 | IM7, rat IgG2b | SuperBright780 | Catalog number: 78-0441-82, RRID:AB_2722968 | ThermoFisher |
| CD45 | 30-F11, rat IgG2b | APC | Catalog number: 103112 RRID:AB_312977 | BioLegend |
| CD45 | 30-F11, rat IgG2b | BB790-P | Special order | BD Biosciences |
| CD45 | 30-F11, rat IgG2b | BUV395 | Catalog number: 564279 RRID:AB_2651134 | BD Biosciences |
| CD45 | 30-F11, rat IgG2b | BUV563 | Catalog number: 565710, RRID:AB_2722550 | BD Biosciences |
| CD45.1 | A20, mouse IgG2a | FITC | Catalog number: 11-0453-82 RRID:AB_465058 | eBioscience |
| CD45.2 | 104, mouse IgG2a | PE | Catalog number: 12-0454-83 RRID:AB_465679 | eBioscience |
| CD45.2 | 104, mouse IgG2a | PerCP-Cy5.5 | Catalog number: 45-0459-42, RRID:AB_10717530 | ThermoFisher |
| CD62L | MEL-14, rat IgG2a | BUV737 | Catalog number: 565213 RRID:AB_2721774 | BD Biosciences |
| CD62L | MEL-14, rat IgG2a | PE-Cy7 | Catalog number: 25-0621-82, RRID:AB_469633 | eBioscience |
| CD64 | X54-5/7.1, mouse IgG1 | AF532 | Catalog number:139302 RRID: AB_10840366 | BioLegend |
| CD64 | X54-5/7.1, mouse IgG1 | PE-Dazzle594 | Catalog number: 139320 RRID:AB_2566559 | BioLegend |
| CD69 | H1.2F3, armenian hamster | PE | Cat# 12-0691-83, RRID:AB_465733 | eBioscience |
| CD69 | H1.2F3, armenian hamster | PE-Cy5 | Catalog number: 15-0691-82 RRID:AB_468772 | eBioscience |
| CD80 | 16-10A1, armenian hamster | BB630 | Special order | BD Biosciences |
| CD8a | 53-6.7, rat IgG2a | APC | Catalog number: 17-0081-83 RRID:AB_469336 | ThermoFisher |
| CD8a | 53-6.7, rat IgG2a | Biotin | Catalog number: 13-0081-82 RRID:AB_466346 | eBioscience |
| CD8a | 53-6.7, rat IgG2a | BUV805 | Catalog number: 564920 RRID:AB_2716856 | BD Biosciences |
| CD8a | 53-6.7, rat IgG2a | BV785 | Catalog number: 100750, RRID:AB_2562610 | BioLegend |
| CD8a | 53-6.7, rat IgG2a | PerCP-eFluor710 | Catalog number: 46-0081-82, RRID:AB_1834433 | eBioscience |
| CD90.1/Thy1.1 | HIS51, mouse IgG2a | Alexa Fluor 488 | Catalog number: 202506, RRID:AB_492882 | BioLegend |
| CD90.2 | 53-2.1, rat IgG2a | BV510 | Catalog number: 105335 RRID:AB_2566587) | BioLegend |
| CD95 | Jo2, armenian hamster | BB660 | Special order | BD Biosciences |
| CXCR5 | L138D7, rat IgG2b | BV421 | Catalog number: 145511 RRID:AB_2562127 | BioLegend |
| Fixable Viability D | NA | eFluor780 | catalog number: 65-0865-14 | eBioscience |
| Eomes | REA116, human IgG1 | APC | Catalog number: 130-102-917 RRID:AB_2651634 | Miltenyi Biotec |
| F4/80 | BM8, rat IgG2a | PE-Cy5 | Catalog number: 123111, RRID:AB_893494 | BioLegend |
| F4/80 | BM8, rat IgG2a | Pacific Orange | Catalog number: MF48030 RRID:AB_2539704 | eBioscience |
| Foxp3 | FJK-16s, rat IgG2a | Alexa Fluor 488 | Catalog number: 53-5773-82, RRID:AB_763537 | eBioscience |
| Foxp3 | REA788, human IgG1 | APC | Catalog number: 130-111-601 RRID:AB_2651768 | Miltenyi Biotec |
| Foxp3 | FJK-16s, rat IgG2a | APC | Catalog number: 17-5773-82, RRID:AB_469457 | eBioscience |
| Foxp3 | FJK-16s, rat IgG2a | eFluor450 | Catalog number: 48-5773-82, RRID:AB_1518812 | eBioscience |
| Foxp3 | FJK-16s, rat IgG2a | PE-Cy7 | Catalog number: 25-5773-82 RRID:AB_891552 | eBioscience |
| FR4 | 12A4, rat IgG1 | BUV496 | Catalog number: 750415 | BD Biosciences |
| GATA-3 | L50-823, mouse IgG1 | BUV737 | Special order | BD Biosciences |
| GATA-3 | L50-823, mouse IgG1 | PE-Cy7 | Catalog number: 25-9966-42, RRID:AB_2573568 | eBioscience |

| | | | | |
|---|---|---|---|---|
| GFP | FM264G | AF488 | Catalog number: 338008 RRID:AB_2563288 | BioLegend |
| GITR | DTA-1, rat IgG2b | BUV805 | Catalog number:742061 RRID:AB_2871349 | BD Biosciences |
| Helios | 22F6, armenian hamster | e450 | Catalog number: 48-9883-41RRID:AB_2574137 | eBioscience |
| Helios | 22F6, armenian hamster | PE-Dazzle594 | Catalog number: 137232, RRID:AB_2565797 | BioLegend |
| ICOS | C398.4A, armenian hamster | BV605 | Catalog number: 313538 RRID:AB_2687079 | BioLegend |
| IFNg | XMG1.2, rat IgG1 | BV605 | Catalog number: 505839 RRID:AB_2561438 | BioLegend |
| IgD | 11-26c.2a, rat IgG2a | BUV496 | Catalog number: 624283 | BD Biosciences |
| IgE | R35-72, rat IgG1 | BV605 | Catalog number: 744281 RRID:AB_2742118 | BD Biosciences |
| IgM | Il/41, rat IgG2a | PE-Cy5 | Catalog number: 15-5790-82 RRID:AB_494222 | eBioscience |
| IL-2 | JES6-5H4, rat IgG2b | PE | Catalog number: 12-7021-82 RRID:AB_466150 | eBioscience |
| IL-5 | TRFK5, rat IgG1 | PE | Catalog number: 504303, RRID:AB_315327 | BioLegend |
| IRF4 | 3E4, rat IgG1 | AF647 | Discontinued | eBioscience |
| Ki-67 | 16A8, rat IgG2a | AF700 | Catalog number: 652420 RRID:AB_2564285 | BioLegend |
| KLRG1 | 2F1/KLRG1, syrian hamster | BV785 | Catalog number: 138429 RRID:AB_2629749 | BioLegend |
| Ly6C | AL-21, rat IgM | BB790-P | Special order | BD Biosciences |
| Ly6C | HK1.4, rat IgG2c | BV711 | Catalog number: 128037 RRID:AB_2562630 | BioLegend |
| Ly6G | 1A8, rat IgG2a | BV510 | Catalog number: 127633 RRID:AB_2562937 | BioLegend |
| Ly6G | 1A8, rat IgG2a | BV421 | Catalog number:127627 RRID:AB_10897944 | BioLegend |
| Ly6G | 1A8, rat IgG2a | BV650 | Catalog number: 127641 | BioLegend |
| MHCII (I-A/I-E) | M5/114.15.2, rat IgG2b | PerCP | Catalog number: 107624 RRID:AB_2191073 | BioLegend |
| MHCII (I-A/I-E) | M5/114.15.2, rat IgG2b | AF700 | Catalog number: 56-5321-82 RRID:AB_494009 | eBioscience |
| Neuropilin (CD30 | 3DS304M, rat IgG2a | PerCP-eFluor710 | Catalog number: 46-3041-82 RRID:AB_2573741 | eBioscience |
| NK1.1 | PK136, mouse IgG2a | BUV563 | Catalog number: 741233 RRID:AB_2870785 | BD Biosciences |
| NK1.1 | PK136, mouse IgG2a | Biotin | Catalog number: 108702 RRID:AB_313389 | BioLegend |
| NKp46 | 29A1.4, rat IgG2a | FITC | Catalog number:137606 RRID:AB_2298210 | BioLegend |
| PD-1 | 29F.1A12, rat IgG2a | BV711 | Catalog number: 135231 RRID:AB_2566158 | BioLegend |
| PDCA-1 | 927, rat IgG2b | BB700 | Catalog number: 747601, RRID:AB_2744169 | BD Biosciences |
| PDCA-1 | 927, rat IgG2b | BUV737 | Catatalog number: 749272, RRID:AB_2873649 | BD Biosciences |
| PDCA-1 | 927, rat IgG2b | PerCP-Cy5.5 | Catalog number:127022 RRID:AB_2566647 | BioLegend |
| pSTAT3 | 13A3-1, mouse IgG1 | AF647 | Catalog number: 651008, RRID:AB_2572086 | BioLegend |
| pSTAT5 | SRBCZX, mouse IgG1 | PE-eFluor610 | Catalog number: 61-9010-42, RRID:AB_2574672 | eBioscience |
| RORgT | AFKJS-9, rat IgG2a | APC | Catalog number: 17-6988-82 RRID:AB_10609207 | eBioscience |
| RORgT | Q31-378, mouse IgG2a | BV650 | Catalog number: 564723 RRID:AB_2738916 | BD Biosciences |
| Siglec F | E50-2440, rat IgG2a | BUV615P | Special order | BD Biosciences |
| Siglec F | E50-2440, rat IgG2a | PE | Catalog number: 552126 RRID:AB_394341 | BD Biosciences |
| ST2 (IL-33R) | RMST2-2, rat IgG2a | PE-Cy7 | Catalog number: 25-9335-80 RRID:AB_2637463 | eBioscience |
| Streptavidin | NA | AF350 | Catalog number: S11249 | Invitrogen |
| Streptavidin | NA | PE | Catalog number: 12-4317-87 | Invitrogen |
| Streptavidin | NA | Qdot545 | Catalog number: Q10091MP | Invitrogen |
| T-bet | 4B10, mouse IgG1 | Alexa Fluor 594 | Catalog number:644833 RRID:AB_2728473 | BioLegend |
| T-bet | Violet: BV421 & e450 | BV421 | Catalog number: 644832 RRID:AB_2686976 | BioLegend |
| TCR gamma/delta | GL3, armenian hamster | Biotin | Catalog number: 13-5711-85, RRID:AB_466669 | eBioscience |
| TCR gamma/delta | GL3, armenian hamster | FITC | Catalog number: 11-9959-42, RRID:AB_10669049 | eBioscience |
| TCR gamma/delta | GL3, armenian hamster | SuperBright780 | Catalog number: 78-5711-82, RRID:AB_2744919 | eBioscience |
| TCR-beta | H57-597, armenian hamster | BB790 | Special order | BD Biosciences |
| TCR-beta | H57-597, armenian hamster | eFluor450 | Catalog number: 48-5961-82, RRID:AB_11039532 | eBioscience |
| TNFRII | TR75-89, armenian hamster IgG | PE | Catalog number: 113406 RRID:AB_2206941 | BioLegend |
| XCR1 | ZET mouse IgG2b | BV650 | Catalog number: 148220 RRID:AB_2566410 | BioLegend |
| PE/Cy5.5® Conju | NA | PE-Cy5.5 | Catalog number: ab102899 | Abcam |
